# Early Life Shift in Fear Extinction is Driven by Prefrontal-Amygdala Engagement

**DOI:** 10.64898/2026.11.19.767882

**Authors:** Jennifer Wilkin, Anna Elisabete Canella, Abdessattar Khlaifia, Arely Cruz-Sanchez, Hanista Premachandran, Mehreen Inayat, Meltem Nacakli, Samuel Varghese, Maithe Arruda-Carvalho

## Abstract

The medial prefrontal cortex (mPFC) displays a delayed maturation long hypothesized to guide developmental changes in threat processing. Nevertheless, the mechanism by which ontogenetic changes in mPFC function guide the developmental emergence of behavior is unclear. Early life marks a shift between a permanent extinction phenotype typical of juveniles, toward the standard, more transient extinction seen in adult rodents. This shift coincides with developmental changes in mPFC-amygdala connectivity, a pathway necessary for extinction processing in adults. Here we used optogenetics and slice electrophysiology to show that, in contrast to adult animals, juvenile mice displaying permanent extinction fail to engage the mPFC-amygdala pathway at the synaptic, cell activity or behavioral levels. Consistent with a requirement of this pathway for the developmental transition in extinction types, optic potentiation of the mPFC-amygdala pathway accelerates the onset of adult extinction. Our findings show that the developmental onboarding of the mPFC-amygdala pathway drives a shift in extinction processing toward increased fear reemergence that persists throughout adulthood.

## Introduction

Human imaging, postmortem and preclinical work show that brain development follows a pattern of graded changes in anatomical and functional connectivity, with the maturation of brain regions involved in motor and sensory abilities preceding that of regions implicated in top-down control of thoughts and action^1,2^. These dynamic changes are believed to reflect experience-driven fine-tuning of neural circuits toward optimizing behavior^1,3^, but causal links between developmental changes in brain connectivity and age-dependent changes in behavior remain elusive. Threat or fear associations are robust and last throughout the lifetime^4,5^. While fear extinction can suppress fear responses, these often re-emerge with time^6–8^. In contrast, extinction in young rodents is permanent^8–10^, marking a developmental switch around juvenility whereby animals shift from an *immature*-type, permanent extinction toward the *mature*-type, impermanent extinction that is typical of adult animals^8–14^.

The medial prefrontal cortex (mPFC) modulates fear extinction^15–20^, particularly via its projections to the basolateral amygdala (BLA)^21–24^ and exhibits a relatively delayed maturation known to influence emotional regulation^25^. Remarkably, the timing of the extinction switch coincides with changes in mPFC and BLA inhibitory transmission^26–30^ and mPFC-BLA circuit connectivity^31–37^, suggesting that developmental changes in extinction processing may be driven in part by mPFC-BLA maturation. Despite some evidence of differential recruitment of mPFC to fear acquisition and extinction in infancy^10,38,39^, it remains unclear how the delayed maturation of the mPFC and its integration into the threat processing circuitry causally impact fear expression and extinction in early life. To test this, here we examined when and how the developmental engagement of the mPFC-BLA pathway alters fear extinction processing. We found that mPFC-BLA inhibition during extinction disrupts mature-type (but not immature-type) extinction, while mPFC-BLA optic potentiation is sufficient to induce the mature-type extinction phenotype. Our data suggest that the switch in extinction modes occurring in early life is driven by the developmental recruitment of the mPFC-BLA pathway to extinction learning.

## Results

To first establish a precise timeline of the onset of mature-type extinction, we compared male and female C57BL/6J (Jax) mice that underwent auditory fear conditioning at either postnatal day (P)15, P21, P25 or P30. Mice underwent extinction training for 2 days following fear acquisition and were tested for spontaneous recovery a week later (Figure 1A). Younger mice showed higher freezing at fear acquisition (Figure 1B-C), but there were no sex or age differences during extinction (Figure 1D-G). One week later, at spontaneous recovery, P15 animals of both sexes froze less than older animals (Figure 1H-I), consistent with infantile amnesia^40–44^. At P25 and P30 mice showed the return of fear typical of mature-type extinction (Figure 1I), but while P21males showed mature-type extinction, P21 females displayed suppressed freezing consistent with immature-type extinction (Figure 1I). Importantly, this immature-type extinction phenotype was not driven by forgetting, as P21 females retained the tone-shock association in the absence of extinction (Figure S1). This late onset of mature-type extinction in female C57BL/6J mice contrasts with reports of precocious maturation of fear behavior in female rats compared to males^45–47^, a discrepancy that may be driven by differences in retention delay and/or species.

**Figure 1.**
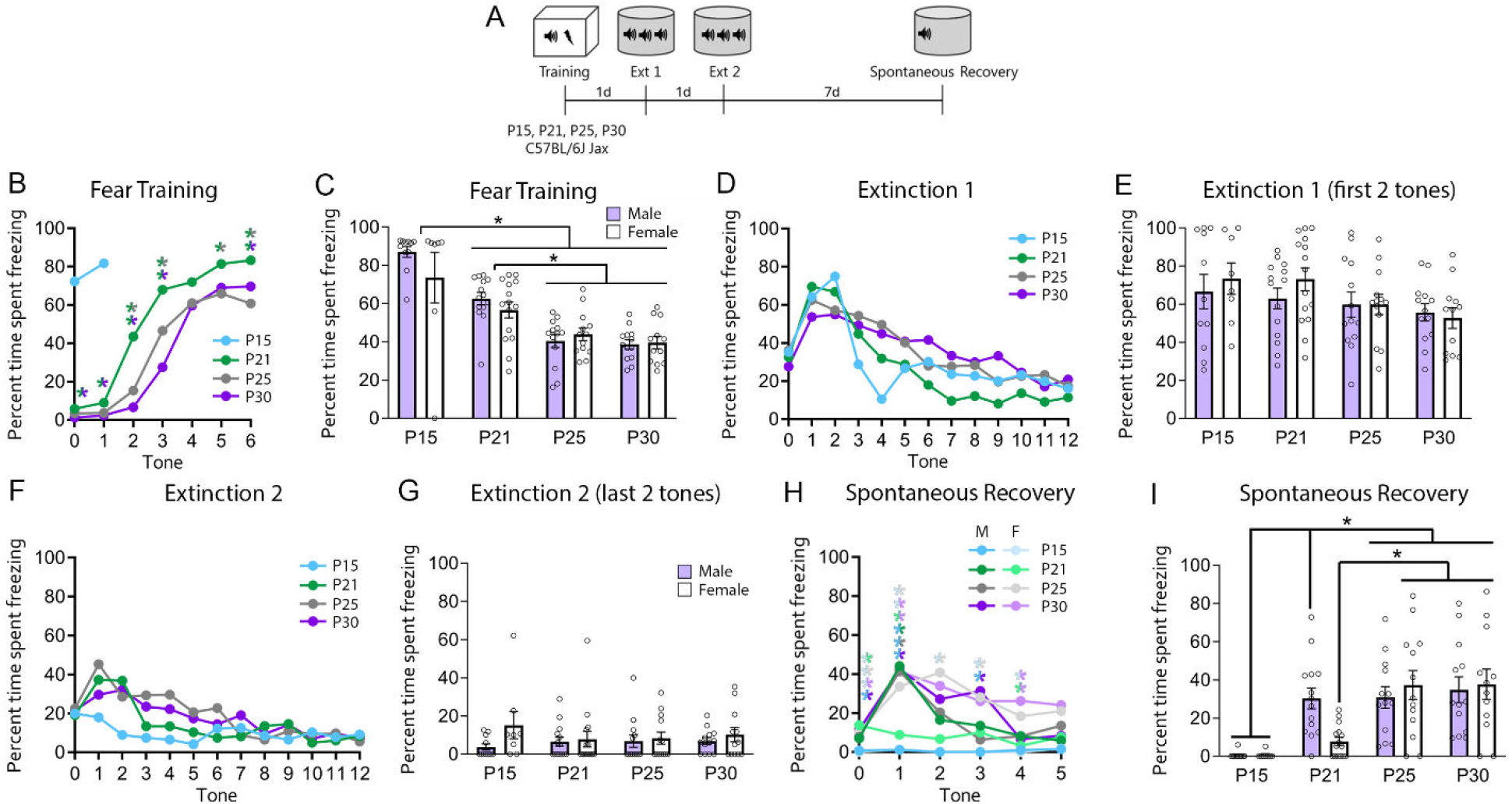
Developmental timeline of the extinction switch in C57BL/6J (Jax) mice. **A**. Experimental timeline. Mice were fear conditioned at different ages, underwent two subsequent days of extinction training, then were tested for spontaneous recovery one week later. **B.** During fear training, freezing increased with each tone in all ages that received six tone-shock pairings (Two-way RM ANOVA with Tukey’s multiple comparisons test: significant effect of tone F_4.418,340.2 =_ 171.7, p < 0.0001, age F_2,77 =_ 21.72, p < 0.0001 and tone x age interaction F_8.835,340.2 =_ 3.965, p < 0.0001). **C.** Average freezing to the tone. P15 and P21 mice exhibited higher average freezing during fear training than P25 and P30 mice (Two-way ANOVA with Tukey’s multiple comparison test: significant effect of age F_3,90 =_ 33.04, p < 0.0001). There were no group differences in freezing across tones (**D;** Three-way RM ANOVA: significant effect of tone F_8.68,789.82 =_ 32.5118, p < 0.0001, age x tone interaction F_26.04,789.82 =_ 2.2730, p = 0.003 but posthocs show p>0.05) or during the first two tones of extinction 1 (**E**). All groups decreased their freezing during extinction 2 (**F;** Three-way RM ANOVA: significant effect of tone F_8.60,782.88 =_ 14.2315, p < 0.0001, age x sex interaction F_3,91 =_ 4.3242, p = 0.0068, age x tone interaction F_25.81,782.88 =_ 2.1121, p = 0.0011, and age x sex x tone interaction F_25.81,782.88 =_ 1.6047, p = 0.0296 but posthocs show p>0.05), and showed equivalent fear suppression at the end of extinction 2 (**G**). **H.** During spontaneous recovery, P15 mice showed lower freezing compared to all other ages in tones 1-3, with P21 females displaying lower freezing compared to P30 females in tones 1 and 4 (Three-way mixed ANOVA: significant effect of age F_3,91 =_ 14.2291, p < 0.0001, tone F_4.32,393.53 =_ 19.1763, p < 0.0001, age x tone interaction F_12.97,393.53 =_ 3.0247, p = 0.0003, sex x tone interaction F_4.32,393.53 =_ 3.9139, p = 0.0239 and age x sex x tone interaction F_12.97,393.53 =_ 2.2709, p = 0.0069) **I.** Average freezing during spontaneous recovery was greater in P25, P30 mice and P21 males compared to P15; and P21 females showed lower freezing compared to older females (Two-way ANOVA significant effect of age: F_3, 91 =_ 14.54, p < 0.0001; age x sex interaction F_3, 91 =_ 3.004, p = 0.0345). P15M n = 11, P15F = 8, P21M n = 14, P21F = 15, P25M n = 13, P25F = 13, P30M n = 13, P30F = 12.

We next tested the necessity of the mPFC-BLA pathway to each extinction phenotype by infusing AAV expressing either the inhibitory opsin archaerhodopsin-3 (ArchT; AAV1-CAG-ArchT-GFP) or control GFP (AAV1-CAG-GFP) into the infralimbic (IL) subregion of the mPFC, and implanting optic fibers over the BLA to selectively inhibit mPFC-BLA projections during extinction training in mice fear conditioned at P21 or P30 (Figure 2A; viral and fiber maps in Figure S2). This manipulation impairs subsequent extinction retrieval in adult mice^21^, and leads to mPFC inhibition in younger mice (Figure S3). There were no group differences at fear acquisition or extinction training (Figure 2B-F). At extinction retrieval, in the absence of laser stimulation, ArchT P30 males, females and P21 males showed higher freezing compared to GFP controls (Figure 2G-H), indicating that the mPFC-BLA pathway is required for subsequent extinction retrieval in these groups. However, ArchT P21 female mice showed robust fear suppression at extinction retrieval (Figure 2H), suggesting that mPFC-BLA activity is not necessary for extinction processing in P21 females. Consistent with our optogenetic manipulation, P21 females show decreased IL (Figure 2I; Figure S4A-E) and IL-BLA (Figure 2J; Figure S4F-K; see Figure S5 for viral maps) cFos expression following extinction.

**Figure 2.**
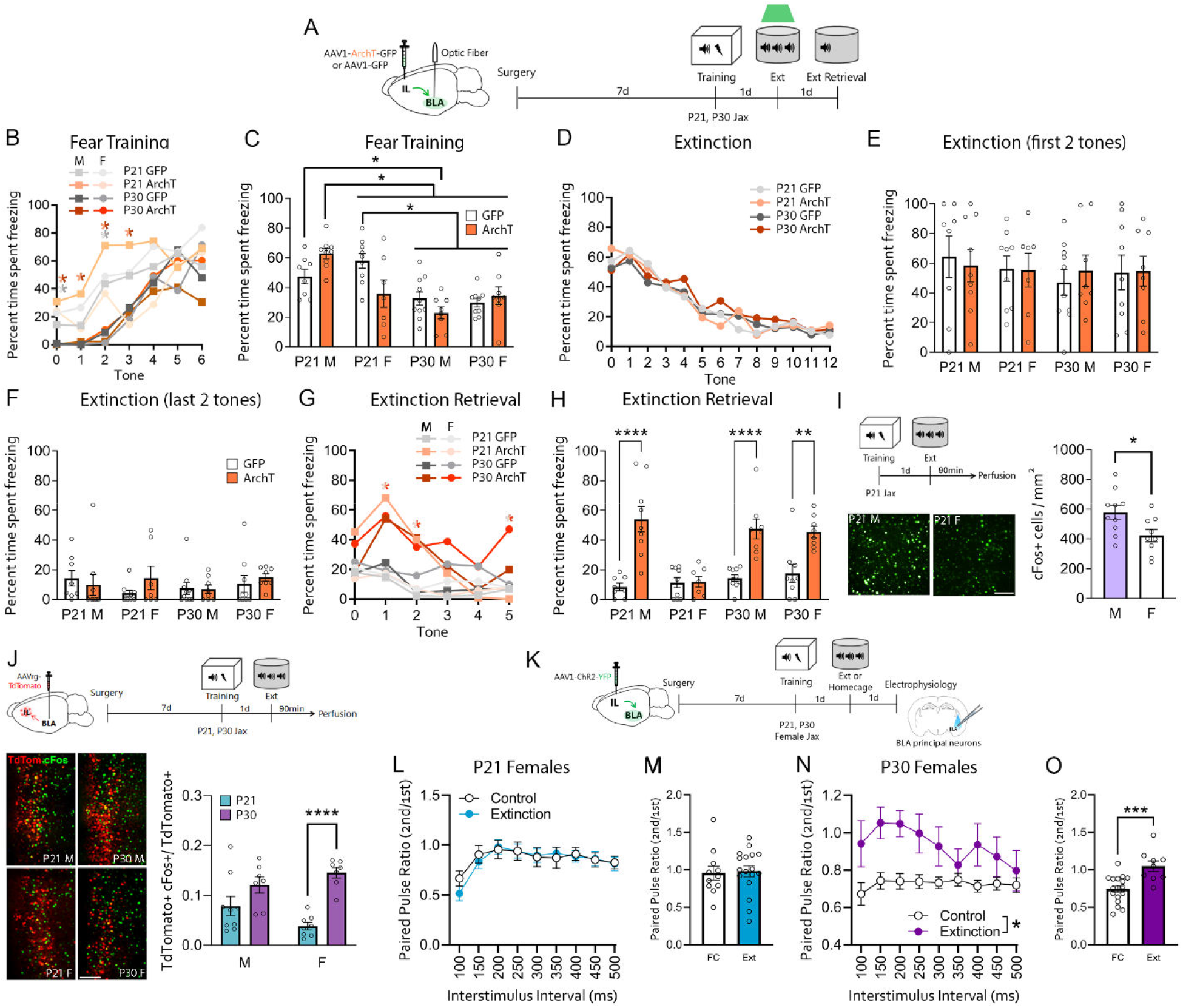
mPFC-BLA recruitment is not necessary for immature-type extinction in C57BL/6J (Jax) mice. **A.** Experimental timeline. Jax mice of different ages were injected with ArchT or GFP constructs and implanted with optic fibers over the BLA one week prior to auditory fear training followed by extinction with laser stimulation to inhibit the mPFC-BLA pathway. Mice were tested the following day in the absence of stimulation. **B.** During fear training, all groups showed increased freezing to each tone, with younger mice showing greater freezing overall (Four-way RM ANOVA: significant effect of age F_1,60 =_ 38.3067, p < 0.0001 and tone F_4.76,285.58 =_ 58.2609, p = < 0.0001, sex x tone interaction F_4.76,285.58 =_ 3.9097, p = 0.0023, age x tone interaction F_4.76,285.58 =_ 4.5502, p = 0.0007, age x virus x sex interaction F_1,60 =_ 11.6069, p = 0.0012, and virus x sex x tone interaction F_4.76,285.58 =_ 2.5418, p = 0.0309). **C**. During fear training, P21 mice generally froze more than P30 mice (Three-way ANOVA with Tukey’s multiple comparisons test: significant effect of age F_1,60 =_ 34.99, p < 0.0001, virus x age x sex interaction F_1,60 =_ 13.33, p = 0.0006). **D.** During extinction, animals showed progressive decrease in freezing across tones (Four-way RM ANOVA: significant effect of tone F_7.51,450.40 =_ 43.9795, p < 0.0001). **E-F.** Freezing to the first two tones (**E**) and last two tones (**F**) of extinction was equivalent across all groups. **G.** At extinction retrieval, tone-by-tone analysis showed that P21 ArchT females had lower freezing than P30 ArchT females at tones 1, 2 and 5 (Four-way RM ANOVA: significant effect of tone F_4.53,271.87 =_ 16.7743, p < 0.0001, age F_1,60 =_ 12.7274, p = 0.0007, virus F_1,60 =_ 24.4160, p < 0.0001, age x sex interaction F_1,60 =_ 10.8078, p = 0.0017, sex x tone interaction F_4.53,271.87 =_ 5.1873, p = 0.003, virus x tone interaction F_4.53,271.87 =_ 7.4551, p < 0.0001, age x sex x virus interaction F_1,60 =_ 7.0335, p = 0.0102, and sex x virus x tone interaction F_4.53,271.87 =_ 3.3204, p = 0.0082). **H.** Average freezing during extinction retrieval. P21 male, P30 male and P30 female ArchT animals froze significantly more than GFP controls, but P21 female GFP and ArchT animals had equivalent freezing (Three-way ANOVA with Tukey’s multiple comparisons test: significant effect of age F_1,60 =_ 7.279, p = 0.0090, sex F_1,60 =_ 6.954, p = 0.0106, virus F_1,60 =_ 54.59, p < 0.0001, virus x sex F_1,60 =_ 11.84, p = 0.0011, age x sex F_1,60 =_ 7.698, p = 0.0074 and virus x age x sex F_1,60 =_ 7.579, p = 0.0078). P21M GFP n = 8, P21F ArchT n = 9, P2F1 GFP n = 9. P21F ArchT = 7, P30M GFP n = 10, 30M ArchT n = 8, P30F GFP n = 9. P30F ArchT = 8. **I.** Extinction-induced cFos activation of IL neurons in P21 male and female mice. *Left, top*: experimental timeline. Mice underwent fear training at P21 followed by one extinction session the next day, and perfusion 90 minutes later. *Left, bottom*: representative images of immunolabeled cFos neurons in IL. *Right:* P21 males showed higher IL cFos+ density than P21 females (unpaired t test, t_16 =_ 2.288, p = 0.0361). M n = 9, F n = 9. **J.** Extinction-induced cFos activation of BLA-projecting IL neurons in P21 and P30 mice. *Top:* Experimental timeline. Mice were infused with a retrograde AAV virus expressing TdTomato into the BLA one week prior to undergoing fear training at P21 or P30. On the next day, mice underwent extinction training and were perfused 90 minutes later. *Bottom* l*eft:* Representative images of co-labeled BLA-projecting (TdTomato+, red) neurons and cFos (green) in IL. *Bottom right:* P21 females show decreased extinction-induced IL-BLA activation compared to P30 females (Two-way ANOVA with Sidak’s multiple comparisons test: significant effect of age F_1,27 =_ 25.40, p < 0.0001 and age x sex interaction F_1,27 =_ 4.738, p = 0.0384). P21M n = 9, P21F = 8, P30M n = 7, P30F = 7. **K-O.** Extinction-driven changes in mPFC-BLA postsynaptic strength in P21 (**L-M**) and P30 (**N-O**) females. **T.** Experimental timeline. Female mice were infused with AAV1-ChR2-YFP virus into the IL and one week later underwent fear training at P21 or P30, followed by extinction (Ext) or remaining in the home cage (fear conditioned only control, FC) the next day. Brains were subsequently processed for slice recordings of BLA principal neurons upon mPFC terminal stimulation with blue light^23^. P21 fear conditioned control and extinguished females showed no difference in paired pulse ratios (**U:** Across interstimulus intervals (ISIs) Two-way RM ANOVA: significant effect of ISI F_3.873, 100.7 =_ 11.14, p < 0.0001; **V:** No group differences at 200ms ISI). In contrast, P30 females showed an extinction-driven increase in mPFC-BLA paired pulse ratio (**W;** Two-way RM ANOVA: significant effect of extinction F_1,225 =_ 46.68, p < 0.0001; **X:** 200ms: unpaired t-test: t_24 =_ 3.869, p = 0.0007), similar to what has been reported in adult animals^23^. * = p < 0.05, *** = p < 0.001, **** = p < 0.0001. Bi-colored asterisks indicate significant differences between groups of the corresponding colors. Full statistical analyses, including non-significant results and post-hocs are included in **Table S1**.

Furthermore, P21 females also show an absence of the extinction-driven mPFC-BLA synaptic plasticity (Figure 2K) seen in P30 females (Figure 2O) and adult mice^23^, reinforcing a lack of recruitment of the mPFC-pathway to extinction processing in mice displaying immature-type extinction. These data suggest a developmental shift in the circuit requirements for extinction, whereby the mPFC-BLA pathway is not necessary for subsequent extinction retrieval in animals displaying immature-type extinction, but is engaged at the synaptic, cellular and behavioral levels in mature-type extinction.

To establish whether mPFC-BLA recruitment is a conserved mechanism marking the transition between extinction profiles, we next tested whether a similar shift in mPFC-BLA engagement would occur in a separate substrain of mice displaying a distinct ontogenetic timeline for fear extinction^48^. C57BL/6NTac (Taconic) P18 and P21 female mice show equivalent fear acquisition (Figure 3A-C) and extinction (Figure 3D-G), but display mature-type extinction earlier than C57BL/6J (Jax) mice, from P21 (Figure 3H-I). Importantly, Taconic females already show memory persistence from P18^48^, confirming that immature-type extinction in Taconic P18 females is not driven by forgetting. We next tested the brain circuit requirements for this behavioral transition. Similar to our previous findings with Jax mice, optogenetic inhibition of the mPFC-BLA during extinction training did not affect extinction retrieval in P18 Taconic mice that display immature-type extinction, but impaired performance in P21 Taconic females that show mature-type extinction (Figure 3J-Q; see Figure S6 for viral and fiber maps). Consistent with a lack of mPFC-BLA pathway recruitment in extinction processing by P18 Taconic females, they also show decreased extinction-driven IL activation compared to P21 females displaying mature-type extinction (Figure 3R, Figure S7). These data suggest that mPFC-BLA engagement is a conserved mechanism marking the switch toward mature-type extinction across mouse substrains.

**Figure 3.**
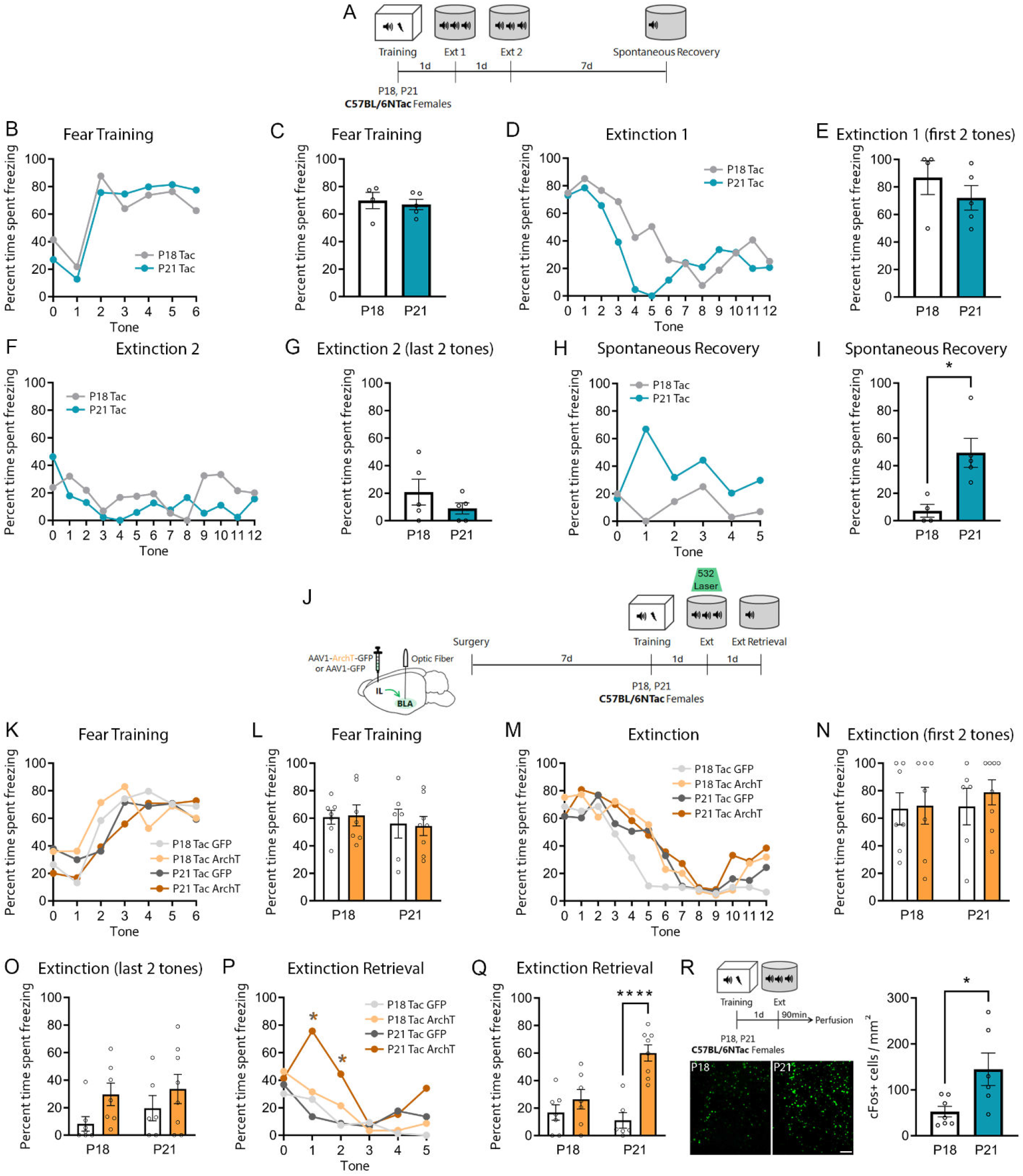
mPFC-BLA engagement marks the transition to mature-type extinction in a mouse substrain that exhibits a distinct ontogenetic fear profile. **A-I**. C57BL6/Tac (Tac) mice extinction switch occurs between P18 and P21 in females. A. Experimental timeline. Female Taconic mice underwent fear training at either P18 or P21, followed by two days of extinction training and spontaneous recovery seven days later. **B.** During fear training, freezing increased across tones in both ages (Two-way RM ANOVA, significant effect of tone F_2.533, 20.27 =_ 11.31, p = 0.002). **C.** There were no age differences in average freezing to the tone during fear training. **D.** During extinction 1, freezing decreased with each tone in both ages (Two-way RM ANOVA, significant effect of tone F_4.643, 37.14 =_ 6.257, p = 0.0003). **E-G**. P18 and P21 Taconic mice did not differ in their freezing rates during the first two tones of extinction 1 (**E**); or across tones (**F**) and at the last two tones (**G**) of extinction 2. **H-I.** During spontaneous recovery, P21 Taconic females showed higher average freezing compared to P18 (**I**; unpaired t-test, t_8 =_ 2.769, p =0.0243). P18 n = 5, P21 n = 5. **J-Q.** Optogenetic inhibition of mPFC-BLA projections during extinction in Taconic female mice. **J.** Experimental timeline. Taconic female mice were injected with ArchT or GFP constructs and implanted with optic fibers over the BLA one week prior to auditory fear training at P18 or P21, followed by extinction in the presence of laser to inhibit the mPFC-BLA pathway. Taconic mice were tested for extinction retrieval the following day in the absence of stimulation. During fear training, all groups showed increased freezing with each tone (**K:** Three-way RM ANOVA: significant effect of tone F_6,144 =_ 19.69, p < 0.0001; **L**). During extinction training, all groups decreased freezing across tones (**M;** Three-way RM ANOVA: significant effect of tone F_5.834, 140 =_ 24.46, p < 0.0001), and had equivalent responses at the start (**N**: first two tones) and end of the session (**O**: last two tones of extinction). mPFC-BLA inhibition during extinction training led to increased freezing at extinction retrieval in P21, but not P18 Taconic mice (**P;** Three-way RM ANOVA with Tukey’s multiple comparisons test significant effect of tone F_3.852, 92.44 =_ 12.35, p < 0.001, age F_1,24 =_ 5.801, p = 0.0241, virus F_1,24 =_ 9.673, p = 0.0048, and interaction F_3.852, 92.44 =_ 3.224, p = 0.0171; **Q:** Two-way ANOVA with Šídák’s multiple comparisons significant effect of age F_1,24 =_ 5.137, p = 0.0327, virus F_1,24 =_ 22.73, p < 0.0001 and interaction F_1,24 =_ 10.18, p = 0.0039). P18 GFP n = 7, P18 ArchT n = 7, P21 GFP n = 6, P21 ArchT n = 8. R. Extinction-induced cFos activation of IL neurons in P18 and P21 female Taconic mice. *Left, top*: Experimental timeline. Taconic mice underwent fear training at either P18 or P21 followed by one extinction session the next day, and perfusion 90 minutes later. *Left, bottom*: Representative images of immunolabeled cFos neurons in IL. *Right:* P21 Taconic females showed higher IL cFos+ density than P18 females (unpaired t test, t_11=_ 2.652, p= 0.0225). P18 n=7, P21 n=6. * = p < 0.05, *** = p < 0.001, **** = p < 0.0001. Bi-colored asterisks indicate significant differences between groups of the corresponding colors. Full statistical analyses, including non-significant results and post-hocs are included in **Table S1**.

To further explore the causal relationship between mPFC-BLA recruitment and mature-type extinction, we next tested whether potentiating mPFC-BLA synapses using an optic long-term potentiation (LTP) protocol^49^ could accelerate the emergence of adult-type extinction in P21 Jax females, which display immature-type extinction (Figure 1). We infused Jax female mice with AAV virus expressing either the excitatory opsin Chronos (pAAV1-Syn-Chronos-GFP) or control GFP into the IL and implanted optic fibers over the BLA (see Figure S8 for viral and fiber maps). At P20, mice underwent an optical LTP protocol in the home cage to potentiate the mPFC-BLA pathway (Figure 4A), leading to the postsynaptic strengthening of mPFC-BLA synapses (Figure S9). On the next day, mice underwent fear conditioning, followed by two days of extinction in the absence of stimulation (Figure 4A). mPFC-BLA LTP did not affect fear acquisition or extinction (Figure 4B-G). In contrast, mPFC-BLA LTP led to increased freezing during spontaneous recovery (Figure 4H-I), suggesting that potentiation of the mPFC-BLA pathway accelerates the onset of mature-type extinction.

**Figure 4.**
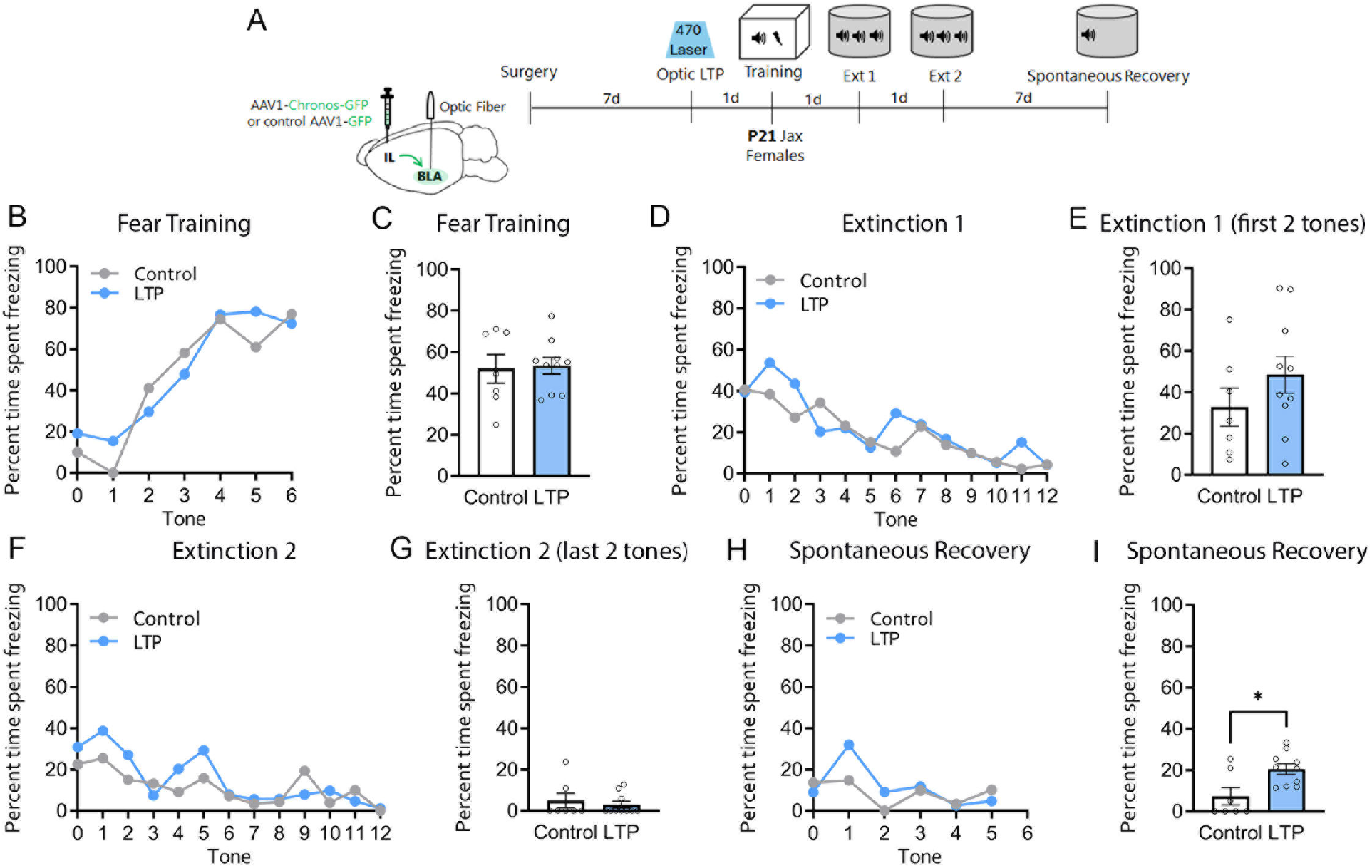
Potentiation of the mPFC-BLA pathway is sufficient to trigger mature-type extinction. **A.** Experimental timeline. Female Jax mice were infused with pAAV1-Syn-Chronos-GFP or AAV1-CAG-GFP into the IL and implanted with optic fibers over the BLA. At P20, all mice underwent an optic LTP protocol in the home cage, and were fear trained the next day (P21), followed by extinction sessions for 2 days, and spontaneous recovery a week afterwards. There were no differences in freezing between control (GFP-infused) and LTP (chronos-infused) animals in fear training (**B**: Tone-by-tone: Two-way RM ANOVA significant effect of tone F_6,90 =_ 19.55, p < 0.0001; **C:** Average tone), extinction 1 (**D**: Tone-by-tone: Two-way RM ANOVA: significant effect of tone F_12,180 =_ 4.566, p < 0.0001; **E**: First two tones) or extinction 2 (**F**: Tone-by-tone: Two-way RM ANOVA: Significant effect of tone F_12,180 =_ 3.719; **G**: Last two tones). During spontaneous recovery, LTP mice froze significantly more than controls (**H:** Tone-by-tone; **I:** Average tone: Unpaired t-test; t_15 =_ 2.874, p = 0.0116). Control n = 7, LTP n = 10. * = p < 0.05. Full statistical analyses, including non-significant results and post-hocs are included in Table S1.

## Discussion

Our findings suggest that the transition toward the mature, transient type of extinction typical of adult rodents is marked by the developmental recruitment of mPFC-BLA activity during extinction training. This is consistent with studies showing that adult mice require IL^50^ and IL-BLA^21,22^ activity during auditory fear extinction for subsequent extinction retrieval and display increased IL activity during successful extinction acquisition^51^ and retrieval^24^. Our data establish that young mice can effectively extinguish auditory fear in the absence of mPFC-BLA activity, but this ability is developmentally restricted and its timing varies across mouse substrains. Critically, while the onset of mPFC-BLA recruitment to extinction processing correlates with pathway-specific developmental changes in synaptic connectivity, inhibition and activity^26,31–35^, which aspect(s) of mPFC-BLA maturation define the closing of this potential critical period remain unknown.

BLA perineuronal nets (PNNs) modulate adult extinction persistence^9^, and developmental increases in BLA PNN number coincide with the timing of the extinction switch^9^. PNNs preferentially form around PV interneurons and stabilize mature inhibitory function^52,53^, suggesting that developmental changes in BLA inhibition may contribute to this developmental transition. mPFC axons synapse onto BLA interneurons in mice^23,33,54,55^, with further evidence of age-dependent changes in mPFC-driven BLA feedforward inhibition^33^ (but see ^35^). Specifically, while our previous work identified a P30 peak in CamKII-mPFC evoked relative inhibition in the BLA of male C57BL/6J mice^33^, subsequent research using both sexes (and a pan-neuronal viral promoter) suggests that the timing of this maturation may differ for females^35^. As neither study covered the timing/sex of the extinction transition identified here, future experiments exploring this maturation in female C57BL/6 mice should test whether mPFC input modulation of BLA PNN maturation contributes to the developmental shift in extinction processing.

Importantly, we found that potentiation of the mPFC-BLA pathway in P21 Jax females was sufficient to trigger the mature-type extinction phenotype, strengthening the link between mPFC-BLA recruitment and the onset of mature-type extinction. Consistent with this finding, manipulations such as early life stress, which accelerate the maturation of and connectivity between the mPFC and BLA in humans^25,56–62^, monkeys^63^, and rodents^60,64–73^ are accompanied by earlier onset of mature-type extinction^74–77^. Our data add to these seminal findings by reinforcing the idea that the maturation of mPFC-BLA connectivity guides how extinction is processed and expressed, with the onboarding of the mPFC potentially triggering increased fear susceptibility to reemergence.

Overall, these data provide empirical support for the long-standing developmental neuroscience tenet that the protracted maturation of the prefrontal cortex guides changes in behavior during early life. The dynamic changes within mPFC-amygdala connectivity taking place in juvenility and adolescence may allow for integration of external information toward an optimized adult neural network, but may also be accompanied by heightened sensitivity to harmful influences carrying important implications for adult behavior^78^. Given that anxiety and other mental disorders featuring deficits in fear processing^75–77^ display an early onset around adolescence^79^, this work highlights the importance of understanding how developmental changes in circuit function may guide windows of plasticity that can be leveraged toward a deeper understanding of the etiology and early interventions for threat-related disorders^80^.

## Materials and Methods

### Animals

Male and female C57BL/6J (Jackson laboratory; Jax) and C57BL/6NTac (Taconic Biosciences; Taconic) mice were bred in-house and ear-notched at P14 for identification. Animals were on a 12h (07:00 to 19:00) light/dark cycle and given food and water *ad libitum*. Pups were designated P0 at the day of birth. Mice were kept with the dam until weaning at P21 and housed in same-sex groups of 2-4 following weaning. Jax mice were weaned two hours before fear conditioning for those trained at P21. Taconic mice were weaned 1h after fear training at P21 as in^48^. For age-matched experiments, litters were split equally across age groups, with approximately equal numbers of male and female animals in each cohort. All mice were kept on ¼” corn cob bedding (The Andersons, cat#10004079) in cages of 10.5”L x 6.5”W x 5”H with 8-12 grams of white crinkled paper nesting material (The Andersons, cat#10004130) and one red Mouse Igloo (Bio-Serv, cat#K3327) per cage. All experiments were performed under the approval of the Animal Care Committee at the University of Toronto.

### Stereotaxic Surgery

Mice underwent stereotaxic surgery 7 days prior to their target age at fear training. Mice were anesthetized with a mixture of ketamine (100 mg/kg) and xylazine (5mg/kg) and placed in a stereotaxic apparatus as previously described^81^. Tear gel (Alcon) was applied to each eye to prevent dehydration. Mice were then secured in a rodent stereotaxic apparatus (Stoelting) using ear bars. Body temperature was maintained at 37°C throughout surgery with a far infrared homeothermic warming pad with a temperature feedback module (RightTemp® Jr, Kent Scientific). A scalpel was used to make an incision down the midline of the scalp, and the connective tissue was excised. A cordless micro drill (#58610, Stoelting) was used to make craniotomies over the target regions. Animals were infused with virus into IL at the following stereotaxic coordinates: P14: AP 1.4, ML ±0.3, DV −2.3; P23: AP 1.7, ML ±0.3, DV −2.5, and viral infusion or fiber implantation into BLA used the following coordinates: P14: AP −1.0, ML ±3, DV −4.9; P23: AP −1.0, ML ±3.1, DV −5.0 in reference to bregma. Virus was delivered with a 500nL Neuros Syringe (#65457-02, Hamilton Company) attached to a probe holder on the stereotaxic apparatus (#751873, Harvard Apparatus). A total of 150-200nL of virus was injected per hemisphere at a rate of 100nL per minute.

For optogenetic manipulations, AAV1-CAG-ArchT-GFP (2 x 10^12^ vg/mL, UNC Lot#AV6295C)^82^ or a control virus AAV1-CAG-GFP (7 x 10^12^ vg/mL, Addgene #37825-AAV1) was infused into the IL region of mPFC, and optic fibers were surgically implanted bilaterally above the BLA. For long-term potentiation experiments, pAAV-Syn-Chronos-GFP (1 x 10^13^ vg/mL, Addgene #59170-AAV1), or a control virus of AAV1-CAG-GFP (7 x 10^12^ vg/mL, Addgene #37825-AAV1) was infused into the IL region of mPFC, and optic fibers were surgically implanted bilaterally above the BLA. Laser power was calibrated for a light output of 20 mW over terminals. For retrograde labelling of BLA-projecting neurons, mice were infused bilaterally with rgAAV1-CAG-TdTomato (7 x 10^12^ vg/mL, Addgene #59462-AAVrg) into the BLA.

### Behavior

#### Apparatus

Auditory fear training occurred in a stainless-steel fear conditioning apparatus (32cm wide,25.5cm high, 25.5cm deep; catalog #VFC-008, Med Associates) that contained shock-grid floors (36 rods, 2mm diameter), referred to as context A. The fear conditioning apparatus was enclosed within a sound-attenuated chamber (63.5cm wide, 36.8cm, high, 124 74.9cm deep; catalog #NIR-022MD, Med Associates). Context A was cleaned with ethanol before and after each experiment. Extinction retrieval, extinction and spontaneous recovery took place in a modified version of the fear training context, which consisted of the same chamber with flexible white plastic panels covering the walls and floor to conceal the underlying shock-grid floor (context B), and was cleaned with water before and after each experiment. Tones originated from a microphone inset into the wall of each chamber. In all experimental protocols, the inter-trial interval (ITI) between each tone presentation was 1 minute, with a 2-minute acclimation period at the beginning of each experiment.

#### Fear Conditioning

On fear training day, after the two-minute acclimation period, mice received six tone-shock pairings (10-sec 2kHz tone of 80 Db paired with a co-terminating 1-sec 0.6 mA foot shock). To ensure equivalent fear expression across ages, P15 mice received one tone-shock pairing. On days 2 (and sometimes 3, depending on experimental design), animals underwent an extinction session in a different context (context B) consisting of twelve blocks of 10-sec tones (2kHz, 80 Db) in the absence of shock. One week following the second extinction day, mice underwent spontaneous recovery, which consisted of five 10-sec tones (2kHz, 80 Db) in context B.

For the optogenetic inhibition experiments, animals that underwent surgery began habituation 5 days after surgery for 2 consecutive days. For the 2 days of habituation, animals were transported to the fear room where they were handled for 5 minutes by the experimenter, then attached to the patch cord and placed in the fear chamber for 10 minutes each day. Once mice reached their target age, they underwent the same fear conditioning protocol described above. Mice underwent one session of extinction training on the following day, consisting of twelve 10-sec tones in context B (2kHz, 80 Db). During extinction training, 532 nm laser (Laserglow Technologies; LRS-0532 DPSS laser) was shone onto BLA during the presentation of all twelve 10-sec tones (adapted from^21^). Laser power was set to 20mW measured at the tip of the optic fiber. Laser power was calibrated before each experiment by measuring the power at the tip of the patch cord with a PM100D optical power meter with an S120C sensor (Thorlabs). Mice then underwent an extinction retrieval test one day following extinction, consisting of five 10-sec tones (2kHz, 80 Db) in context B in the absence of laser stimulation.

#### Behavioral Analyses

Freezing behavior during the CS was defined as the absence of all movement other than that required for respiration^83,84^. Freezing quantification was automated using MedAssociates VideoFREEZE software (motion threshold 18, lasting longer than 1 second)^55^.

For experiments using optogenetic manipulation, due to interference of the patch cord with automated assessment, freezing was hand-scored by a blind experimenter, and calculated using percent component freezing to each tone. Manual scoring exhibit high similarity (98%) with VideoFreeze automated scores in other experimental cohorts. Due to known sex differences in fear behavioral profiles^85^, we manually scored behavior for darting and grooming (data not shown). No significant darting events were found in any age group or sex, which may be due to the relative small dimension of our fear chambers. We also manually scored freezing for P15 fear training data (not subsequent tests), as the small size of the mice and their ability to hide behind a protruding lip on the chamber floor renders automated scoring unreliable^86^.

Similar to previous studies, we excluded animals that showed high (>85%) baseline freezing^8,87,88^, and/or did not show CS-specific freezing after fear conditioning (i.e., if they did not show an increase in freezing from baseline to the first CS presentation)^89,90^ or any reduction in freezing from the first tone to the last tone of extinction^91^. We applied these criteria to ensure that 1) freezing to the CS presentations was associative and not generalization, 2) the fear memory was acquired, and 3) within-session extinction occurred to make accurate inferences about extinction permanence. Also, due to unanticipated power outages, the fear training data from three P15 animals was corrupted and could not be recovered.

The phenomenon of within-session extinction is often observed in tests consisting of multiple CSs, with a decrease in freezing behavior within a session not necessarily indicative of extinction retention^92^. When analyzing cued freezing in our behavioral tests, the first two tones were used to assess fear retrieval or spontaneous recovery, and the last two to determine extinction efficacy (as done in^9^). Freezing data from each tone of Extinction 2, and average freezing for the end of Extinction 2, was also analyzed for successful extinction learning across age groups.

### Statistical Analysis

Data were analyzed used GraphPad Prism version 11.0.0 for Windows, GraphPad Software, Boston, Massachusetts USA, and R (R Core Team, 2025). Freezing data was compared using unpaired t-tests, two-way ANOVAs, three-way ANOVAs and four-way ANOVAs for repeated measures data. Post-doc comparisons were performed using Tukey’s test for comparing all means, and Šídák’s multiple comparisons test for selected paired comparisons. For clarity and streamlining of statistical reporting, significant ANOVA and t test results are reported in the figures legends, with full statistical analyses (including post-hocs) for all experiments available in **Table S1**. In the absence of sex effects, line graphs were collapsed by sex for visual clarity.

### Optic Long-term Potentiation Protocol

One day prior to fear conditioning, mice were placed in a transfer cage and received 200 seconds of 473 nm blue laser stimulation (Laserglow Technologies; LRS-0473 DPSS laser) independent of viral construct using 20 trains of 150 pulses at 250 Hz, spaced 10 seconds apart (adapted from^49^) and placed back into their home cage immediately afterwards.

### Perfusions and Sectioning

For cFos experiments, mice were euthanized 90 minutes after the end of extinction day 1. Mice were transcardially perfused with 0.1 M phosphate-buffered saline (PBS), followed by cold 4% paraformaldehyde fixative (PFA). The brains were extracted and stored in PFA overnight at 4°C. Brains were sectioned at 50 µm (1 in 5 series) in the coronal plane with a vibratome (Leica, model VT1000). Sections were stored at -20°C in a cryoprotectant solution comprised of 60% glycerol and 0.01% sodium azide in 0.1M PBS^93^.

### Immunohistochemistry

Immunofluorescence staining for cFos was performed as previously described^81^. Sections were rinsed in 0.1M PBS and then incubated in 1% H_2O2 i_n 0.1M PBS to quench endogenous peroxidase activity. Sections were incubated overnight at 4°C with monoclonal rabbit anti cFos (1:8000, Synaptic Systems cat# 226008) diluted in 0.1M Tris-buffered saline and 0.5% (w/v) Roche Blocking Reagent (Sigma-Aldrich, cat# 11096176001). On the following day, sections were rinsed in 0.1M PBS and incubated in donkey anti-rabbit horseradish peroxidase-conjugated secondary antibody (1:500, Jackson ImmunoResearch, USA, cat# 771-036-152).

Tyramide signal amplification was then performed with fluorescein tyramide conjugate (1:400) diluted in 0.1M borate buffer containing 0.01% H_2O2._ Hoescht 3342 (1:1000 diluted in 0.1M PBS, Thermo Fisher Scientific) was used as a nuclear dye. Sections were then mounted onto gelatin-coated slides, air dried, coverslipped with Permafluor anti-fade mounting medium (Fisher Scientific, cat# TA030FM) and stored in a microscope slide box at 4°C until imaging.

### Imaging and cell counts

Sections containing the IL were imaged in a Nikon NiU epifluorescence microscope at 10x magnification. Cell counts for cFos+ and TdTomato+ cells were performed manually by an experimenter blind to experimental groups using ImageJ Fiji software with the Cell Counter plugin^94^. To quantify neuronal activity in BLA-projecting IL neurons, we performed cell counts of cFos+ IL neurons that were co-labelled with retrograde TdTomato, and divided this count by the total number of TdTomato cells in IL to control for viral load.

### Slice Electrophysiology

Mice were deeply anesthetized with isofluorane and brains were extracted in ice-cold sucrose based cutting solution (195.75 nM sucrose, 2.5 nM KCl, 1.25 nM NaH_2P_O_4,_ 25 nM NaHCO_3,_ 1 nM CaCl_2,_ 4 nM MgCl_2,_ 2 nM Na+ pyruvate and 0.4 nM L-Ascorbic acid with 95% O_2 a_nd 5% CO_2)_. The brains were sectioned in the coronally in 300 μm thick slices using a vibratome (VT1200S, Leica) and transferred to a recovery chamber with artificial cerebrospinal fluid (aCSF: 120 nM NaCl, 2.5 nM KCl, 1.25 nM NaH_2P_O_4,_ 25 nM NaHCO_3,_ 2 nM CaCl_2,_ 1 nM MgCl_2,_ 2 nM Na+ pyruvate and 0.4 nM L-Ascorbic acid with 95% O_2 a_nd 5% CO_2)_ at 30°C where they were recovered for at least 1 hour. After recovery, individual slices were transferred to a submersion chamber, perfused at 2 ml/min with aCSF at room temperature and mounted on an upright microscope (Eclipse FN1, Nikon) equipped with water immersion long-working distance objective (40x) and infrared video camera (DAGE IR-2000).

For the ArchT validation experiment, recordings were obtained 7 days after surgery from IL GFP+ pyramidal neurons with borosilicate pipettes (3-8 MΩ) filled with internal solution containing the following (in mM): 126 K D-Gluconate, 10 HEPES, 0.6 EGTA, 5 KCl, 2 MgCl_2,_ 4 MgATP, 0.3 NaGTP, 5 Na-phosphocreatine. Somatic currents pulses in 25pA increments varying between 0 and 150pA were injected for a total of 1200ms duration. The neurons were simultaneously stimulated with using a TTL-pulsed microscope objective-coupled light-emitting diodes (460 nm, ∼1.36 mW/mm2, 400ms pulses; Prizmatix) 400ms after onset of current injected. The somatic amount of current injected was normalized to rheobase so that prior to light stimulation it elicited between 4-6 action potentials. Number of action potentials were quantified in a 400ms interval before and during light presentation. When quantifying the membrane potential change at baseline, we measured the resting membrane potential before light presentation and compared to the peak amplitude of the negative infection during light presentation^33,95^.

For the LTP validation experiment, whole-cell voltage-clamp recordings were obtained 24 hours after LTP protocol from BLA principal neurons using borosilicate glass electrodes (3–5 MΩ). Electrode internal solution was composed of the following (in mM): 130 Cs-methanesulfonate, 10 HEPES, 0.5 EGTA, 8 NaCl, 5 TEA-Cl, 4 Mg-ATP, 1 QX-314, 10 Na-phosphocreatine, and 0.4 Na-GTP. mPFC axon terminals were stimulated using illumination with a digital multimirror device (Polygon400, Mightex) and a TTL-pulsed microscope objective-coupled light-emitting diode (LED, 470 nm, 14 mW/mm2, Mightex). This pulse intensity evoked maximal response amplitudes. For analysis of AMPA/NMDA ratios, we applied 1 μM TTX (#HB1035, HelloBio) and 100 μM 4-AP (#AB120122, Abcam) for more stringent isolation of monosynaptic transmission^33,55^. All stimulation was conducted at 0.1 Hz to avoid inducing synaptic plasticity. AMPA/NMDA ratio was calculated as the peak EPSC at -70 mV divided by the EPSC amplitude at 100ms after current onset at +40 mV^33,55^.

For the mPFC-BLA paired pulse ratios (PPR), whole cell voltage-clamp recordings were obtained 1.5 hours after extinction retrieval from principal neurons from the medial side of the anterior portion of BLA (similar to^23^) using borosilicate glass electrodes (3–8 MΩ). Electrode internal solution was composed of the following (in mM): 130 Cs-methanesulfonate, 10 HEPES, 0.5 EGTA, 8 NaCl, 4 Mg-ATP, 1 QX-314, 10 Na-phosphocreatine, and 0.4 Na-GTP. mPFC axon terminals were stimulated using a TTL-pulsed microscope objective-coupled light-emitting diodes (460 nm, ∼1.36 mW/mm2, 400ms pulses; Prizmatix or CoolLED pE-300 ultra). All stimulation was conducted at 0.1 Hz to avoid inducing synaptic plasticity. mPFC terminals were stimulated twice per trial in increasing 50 ms intervals between 100 ms and 500 ms. Paired pulse ratios were calculated as the second peak EPSC at -70mV divided by the first within a trial^33,55^.

Data were low-pass filtered at 3 kHz and acquired at 10 kHz using Multiclamp 700B and pClamp 10 (Molecular Devices). Series and membrane resistance was continuously monitored, and recordings were discarded when these measurements changed by >20%. Recordings in which series resistance exceeded 25 MΩ were excluded.

## Acknowledgements

This work was supported by grants from the Canadian Institutes of Health Research CIHR (PJT 399790), Natural Science and Engineering Council of Canada (RGPIN-2017-06344; NSERC CGS-D 569380 for JW), SickKids Foundation and CIHR – Institute of Human Development, Child and Youth Health (NI19-1132R), and Human Frontier Science Program Organization (CDA00009/2018 and RGY0072/2019) to MA-C.

## Author contributions

JW, AEC, AK, ACS, HP, MI, MN and SV performed research and analyzed data. JW and MA-C designed research and wrote the paper.

## Competing interests

The authors declare no competing interests.

## Data, code, and materials availability

All data and code needed to evaluate and reproduce the results in the paper are present in the paper and/or the Supplementary Materials. This study did not generate any new materials. Any additional data are available from the corresponding author upon reasonable request.

## Supplemental Figures and Figure legends

**Figure S1.**
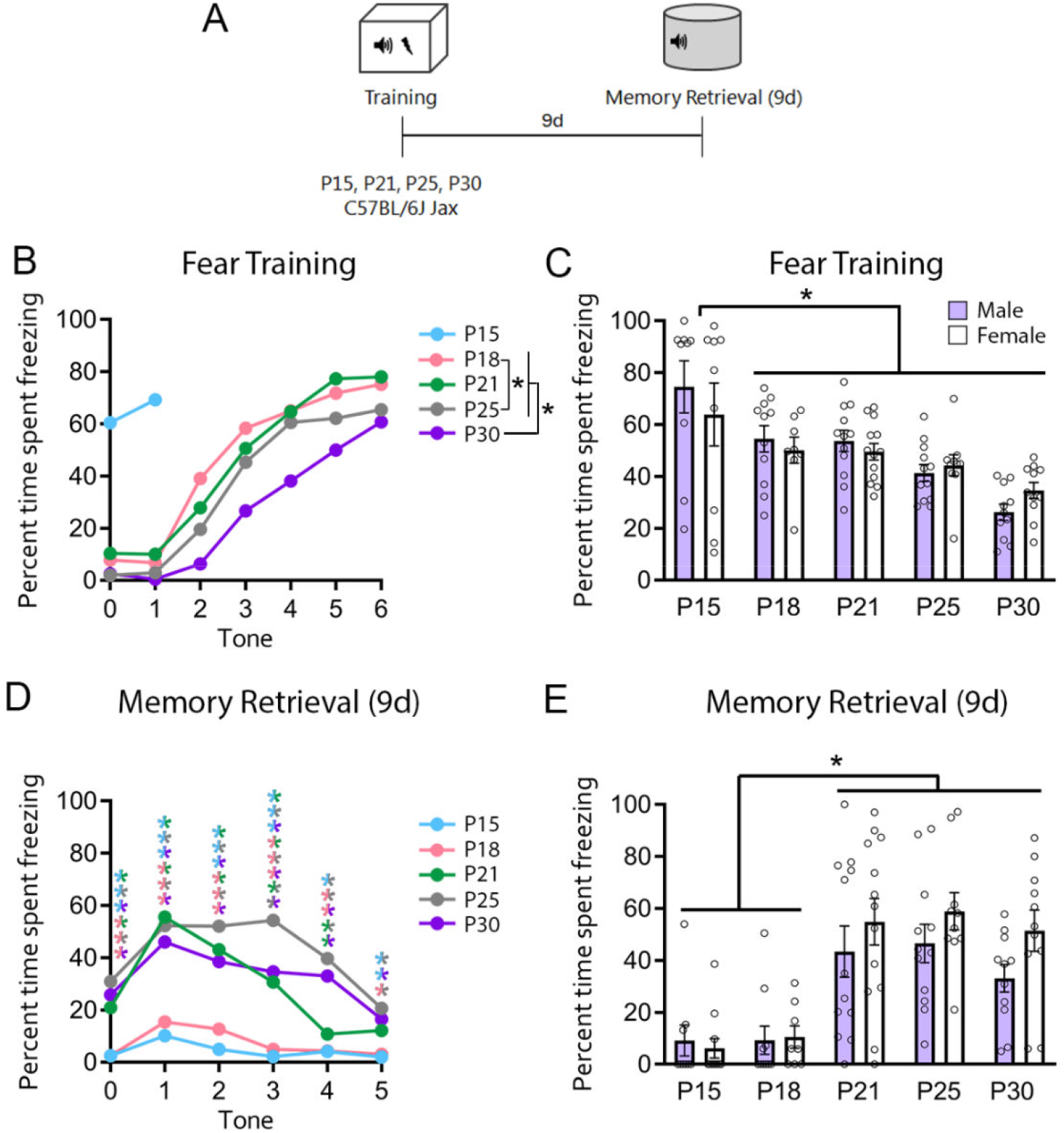
P21 Jax females show memory retention for up to 9 days after fear training. **A.** Experimental timeline. Jax mice were fear trained at different ages, then tested for fear retrieval in the absence of extinction nine days later (same interval as spontaneous recovery in Figure 1A-I). **B-C.** During fear training, tone-by tone analysis showed higher freezing rates in P18 mice compared to P25, with P30 mice displaying lower freezing than younger ages (**B:** Two-way RM ANOVA, significant effect of tone F_4.740, 388.7 =_ 149.9, p < 0.0001 and age F_3,82 =_ 13.67, p < 0.0001), with P15 mice exhibiting higher average freezing than all older ages (**C:** Two-way ANOVA, significant effect of age F_4,96 =_ 12.52, p < 0.0001). **D-E**. During fear retrieval 9 days later, P15 and P18 mice showed significantly lower freezing during the retrieval test than older ages (**D**: Tone-by-tone: Two-way RM ANOVA with Tukey’s multiple comparisons tests, significant effect of tone F_4.143,426.8 =_ 22.27, p < 0.0001, age F_4,103 =_ 19.04, p = 0.0001, and tone x age interaction F_16.57, 426.8 =_ 2.887, p = 0.0001; **E**: Average tone: Two-way ANOVA with Tukey’s multiple comparisons test: significant effect of age F_4,97 =_ 17.15, p < 0.0001). P15M n = 7, P15F = 10, P18M n = 11, P18F = 8, P21M n = 12, P21F = 13, P25M n = 10, P25F = 10, P30M n = 11, P30F = 11. * = p < 0.05, ** = p < 0.01. Bi-colored asterisks indicate significant differences in groups of the corresponding colors. Full statistical analyses, including non-significant results and post-hocs are included in **Table S1**.

**Figure S2.**
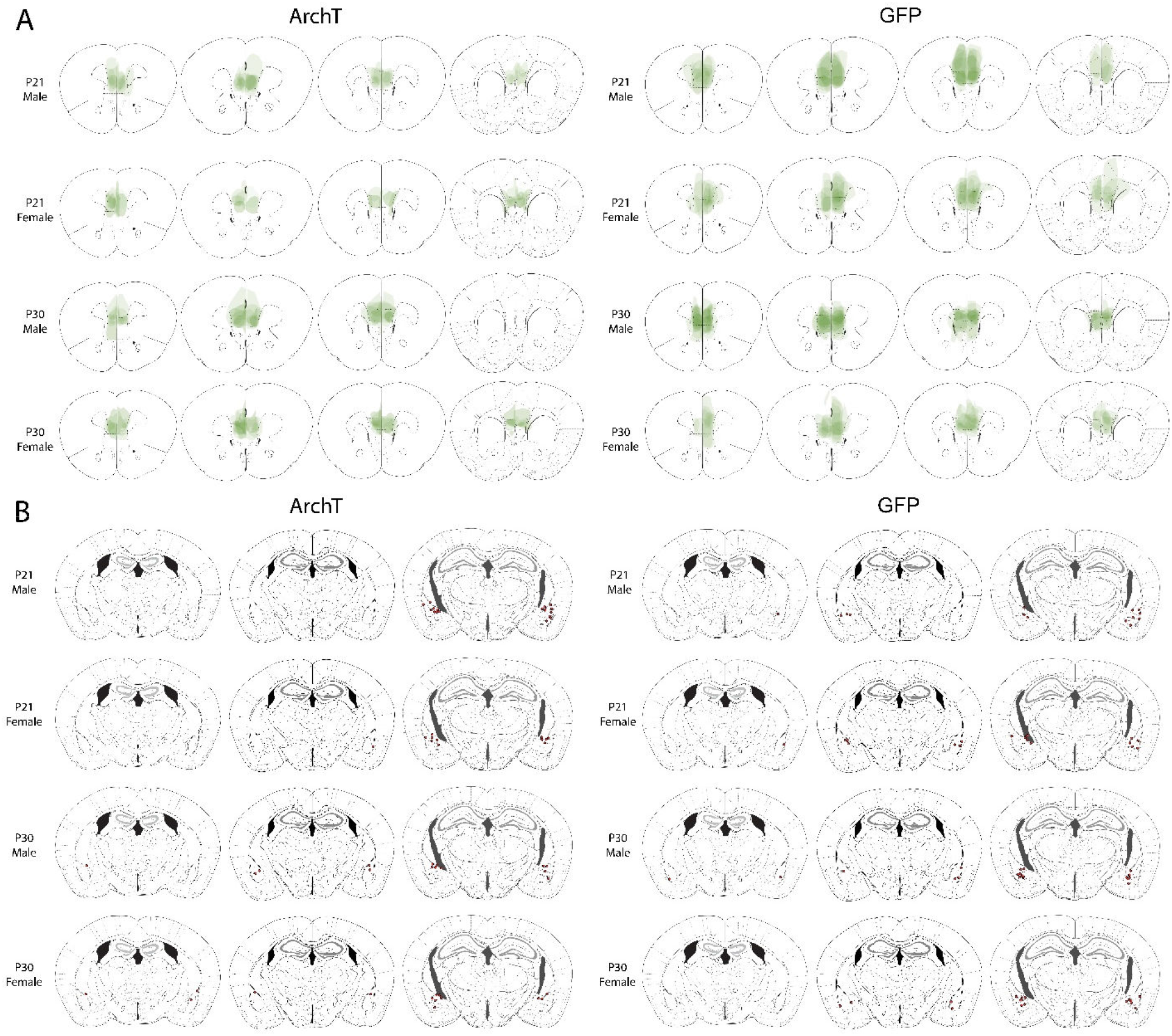
ArchT viral and BLA fiber placement maps for Jax mouse experiments (Figure 1). **A.** Representative diagrams of juxtaposed viral expression in Jax mice injected with ArchT (left) and GFP (right) across groups. **B.** Representative diagrams of BLA fiber placement in mice injected with ArchT (left) and GFP (right) across groups.

**Figure S3.**
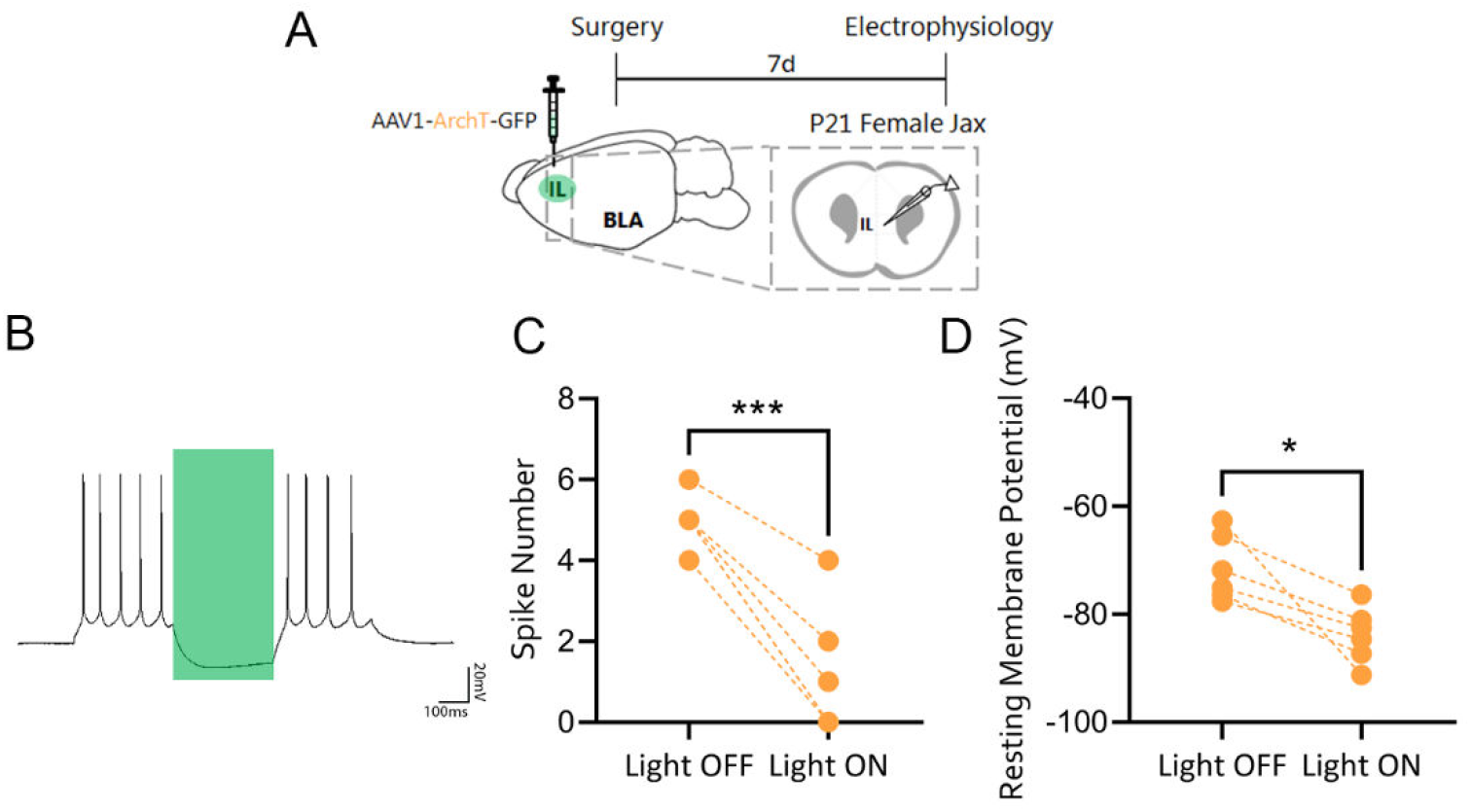
Electrophysiological validation of ArchT experiments. **A**. Experimental timeline. Jax female mice were infused with ArchT AAV in the IL one week prior to patch clamping of IL pyramidal neurons at P21. **B**. Representative trace of action potentials recorded from an ArchT+ neuron in the IL before, during and after 532nm laser stimulation. **C**. The number of action potentials evoked by somatic current injection (50pA) in ArchT+ transfected neurons was significantly decreased following laser stimulation (paired test, t_5 =_ 8.032, p = 0.0005). **D**. The resting membrane potential of ArchT+ neurons became significantly more hyperpolarized following laser stimulation (paired test, t_5 =_ 3.728, p = 0.0136). (n=6 cells, 2 mice). *p<0.05, ***p<0.001.

**Figure S4.**
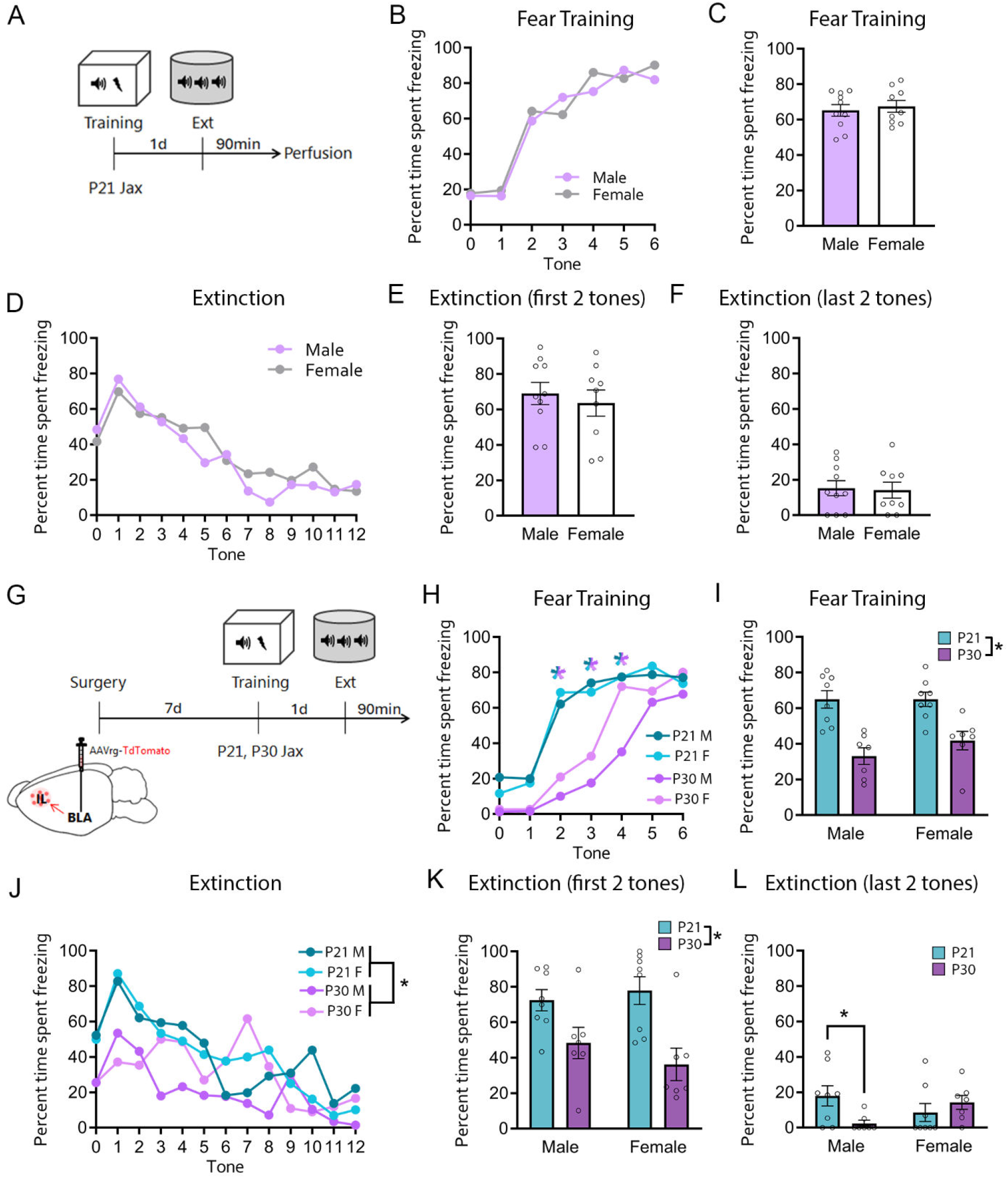
Behavioral data for Jax cFos experiments. Fear conditioning data for the cFos quantifications depicted Figures 1R (**A-F**) and 1S (**G-L**). **A.** Experimental timeline as shown in Figure 1R. P21 male and female Jax mice underwent fear training at P21, followed by one extinction session the following day and perfusion 90 minutes afterwards. **B-C.** There were no group differences during fear training (**B**: Tone-by-tone: Two-way RM ANOVA, significant effect of tone F_6,102 =_ 36.45, p < 0.0001; **C:** Average tone). **D-F.** During extinction, there were no group differences in tone-by tone freezing (**D:** Two-way RM ANOVA: significant effect of tone F_12,204 =_ 8.751, p < 0.0001), during the first two tones (**E**) or last two tones (**F**). M n = 10, F n = 9. **G.** Experimental timeline as shown in Figure 1S. Mice were infused bilaterally in the BLA with AAVrg-TdTomato for retrograde labeling of BLA-projecting IL neurons. Mice were perfused after fear extinction and brain sections underwent cFos immunohistochemistry. **H-I.** During fear training, P21 mice froze more than P30 irrespective of sex (**H:** Tone-by-tone, effect at tones 2, 3 and 4: Three-way RM ANOVA with Sidak’s multiple comparisons test: significant effect of tone F_6,156 =_ 56.18, p < 0.0001, age F_1,26 =_ 31.24, p < 0.0001, tone x age interaction F_6,156 =_ 5.777, p < 0.0001; **I**: Average tone: Two-way ANOVA: significant effect of age F_1,26 =_ 33.86, p < 0.0001). **J-L.** During extinction, P21 mice froze more than P30 males (**J**: Tone-by-tone: Three-way RM ANOVA: significant effect of tone F_12,312 =_ 9.078, p < 0.0001 and age F_1,26 =_ 10.18, p = 0.0037; **K:** First two tones of extinction: Two-way ANOVA: significant effect of age F_1,26 =_ 17.18, p = 0.0003). During the last two tones of extinction, male P21 mice froze more than male P30 mice (**L:** Two-way ANOVA with Sidak’s multiple comparisons test: significant age x sex interaction F_1,26 =_ 5.368, p = 0.0286). P21M n = 8, P21F = 8, P30M n = 7, P30F = 7. *p<0.05. Colored asterisks in Figure 1H indicate group differences between P21 and P30 mice collapsed across sexes. Full statistical analyses, including non-significant results and post-hocs are included in **Table S1**.

**Figure S5.**
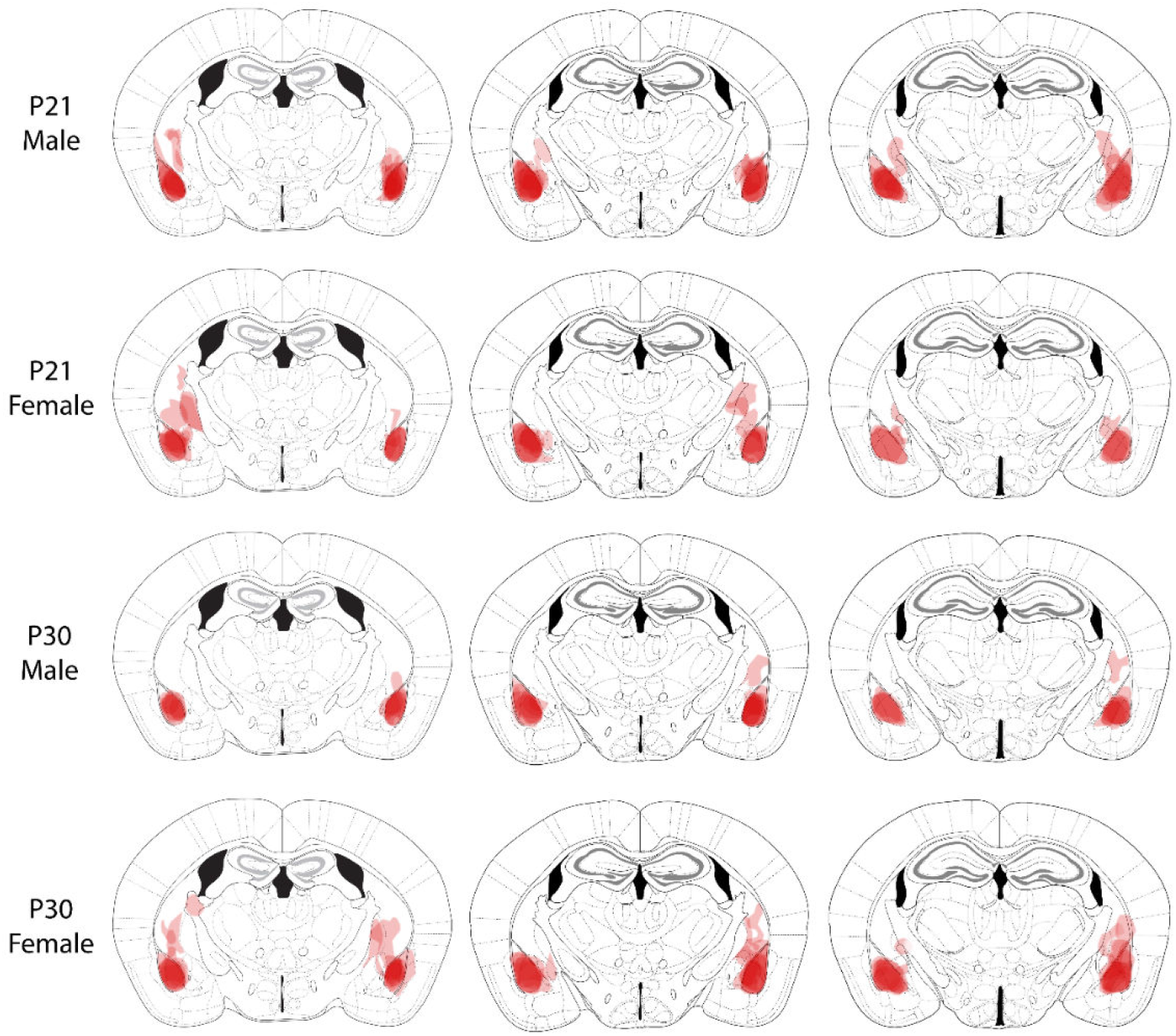
Viral targeting maps for BLA retrograde AAV-TdTomato infusions (Figure 1S). Representative diagrams of juxtaposed viral expression in mice injected with AAVrg-TdTomato in the BLA across groups.

**Figure S6.**
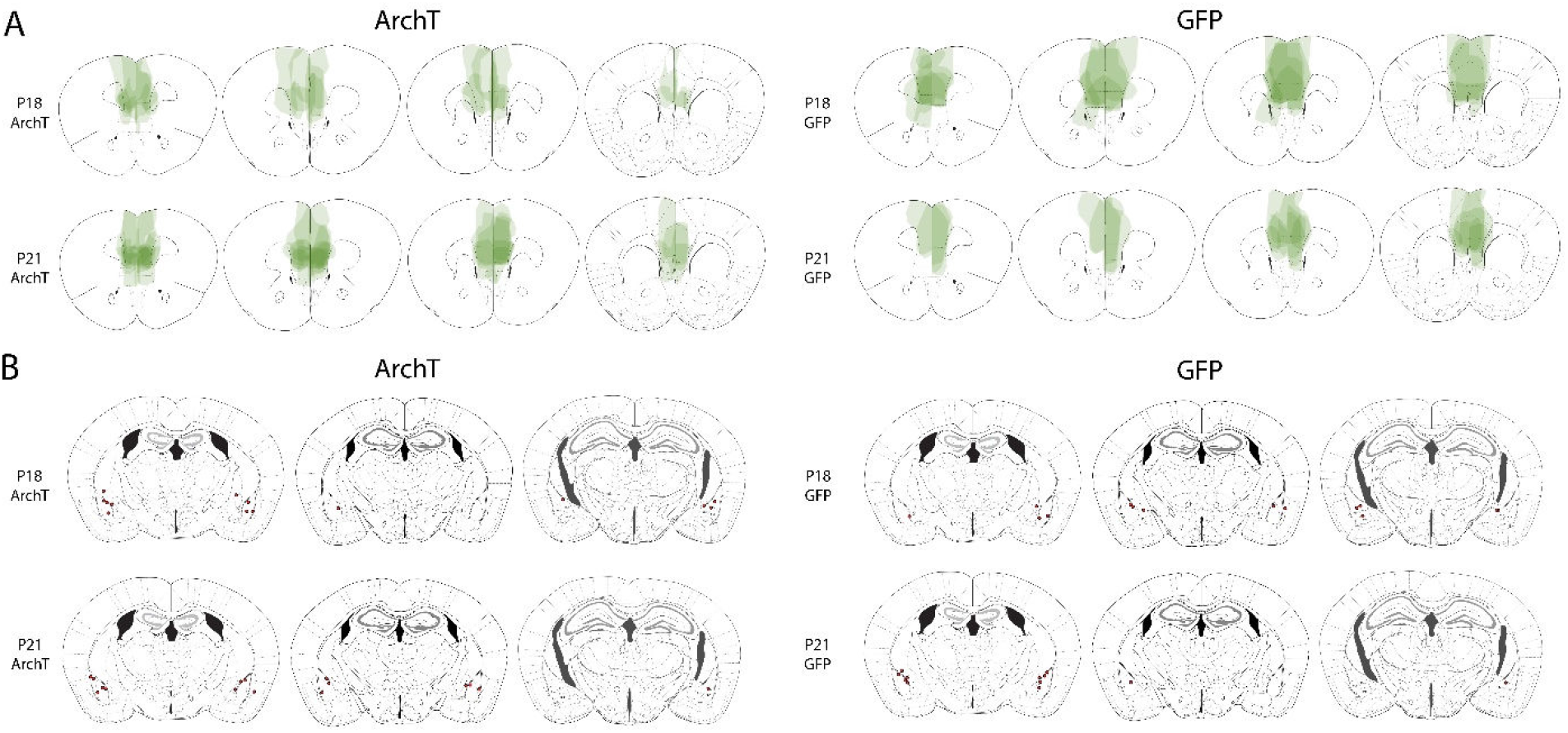
ArchT viral and BLA fiber placement maps for Taconic mouse experiments (Figure 2). **A.** Representative diagrams of juxtaposed viral expression in P18 and P21 female Taconic mice injected with ArchT (left) and GFP (right). **B.** Representative diagrams of BLA fiber placement in Taconic mice injected with ArchT (left) and GFP (right) across groups.

**Figure S7.**
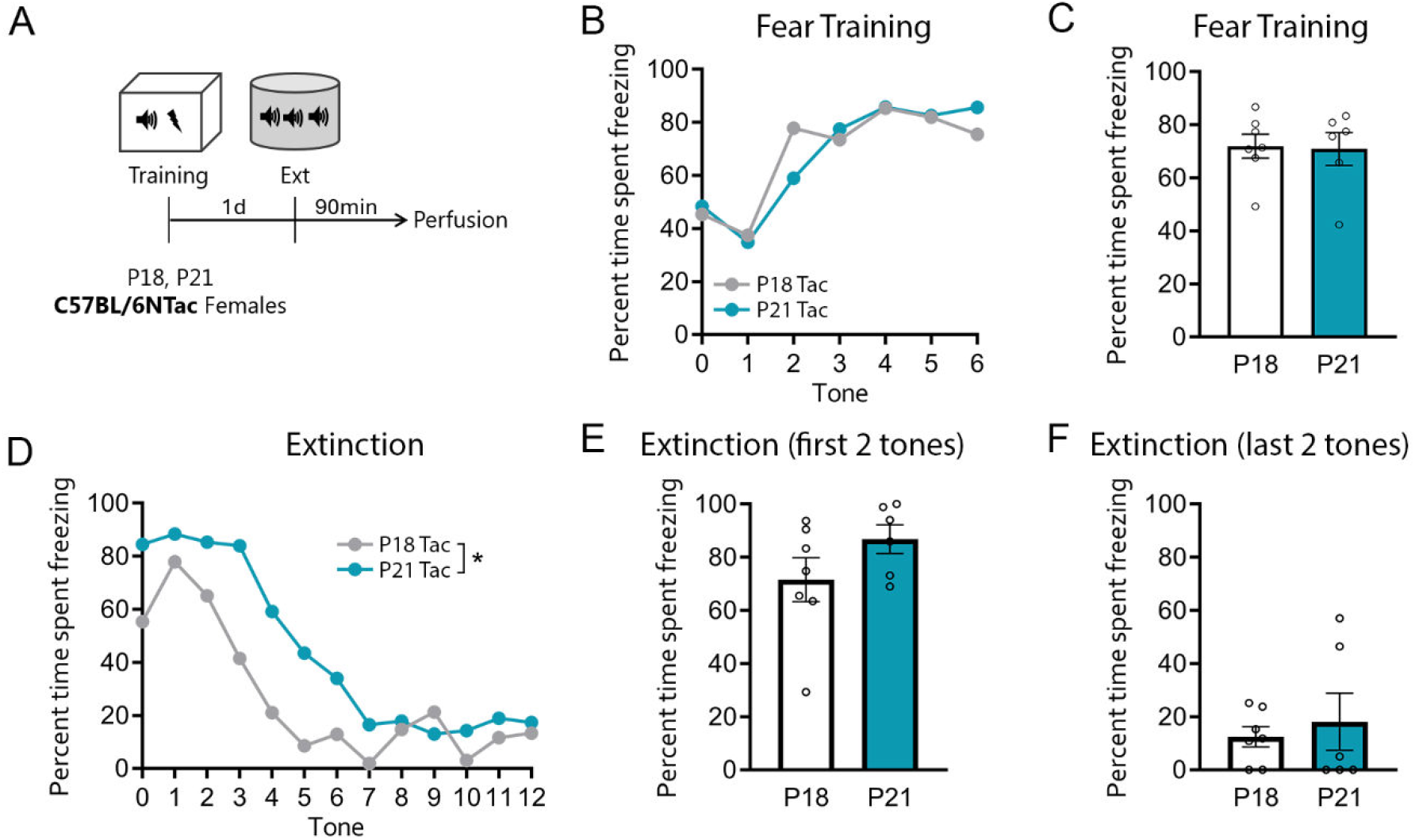
Behavioral data for Taconic cFos experiments. Fear conditioning data for the cFos quantifications depicted Figure 2R. **A.** Experimental timeline as shown in Figure 2R. P18 and P21 female Taconic mice underwent fear training, followed by one extinction session the next day, then were perfused 90 minutes afterwards. **B-C.** During fear training, there were no age differences in tone-by-tone (**B:** Two-way RM ANOVA: significant effect of tone F_3.216, 35.37 =_ 8.262, p = 0.0002) or average freezing (**C**). **D-F.** During extinction, tone-by-tone analysis showed improved within-session extinction in P18 Taconic mice (**D:** Two-way RM ANOVA: significant effect of age F_1,11 =_ 6.179, p = 0.0303 and tone F_5.667, 62.34 =_ 18.74, p < 0.0001), but no group differences during the first two tones (**E**) or last two tones (**F**) of extinction. P18 n = 7, P21 n = 6.

**Figure S8.**
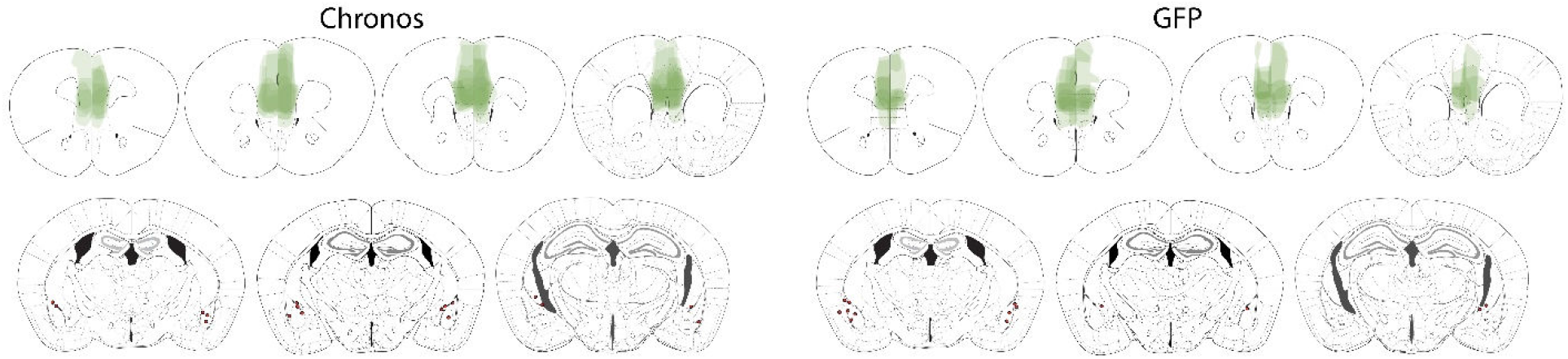
Chronos viral and BLA fiber placement maps for LTP experiment (Figure 3). *Top:* Representative diagrams of juxtaposed viral expression in P21 female Jax mice injected with Chronos (left) and GFP (right). *Bottom:* Representative diagrams of BLA fiber placement in mice injected with Chronos (left) and GFP (right).

**Figure S9.**
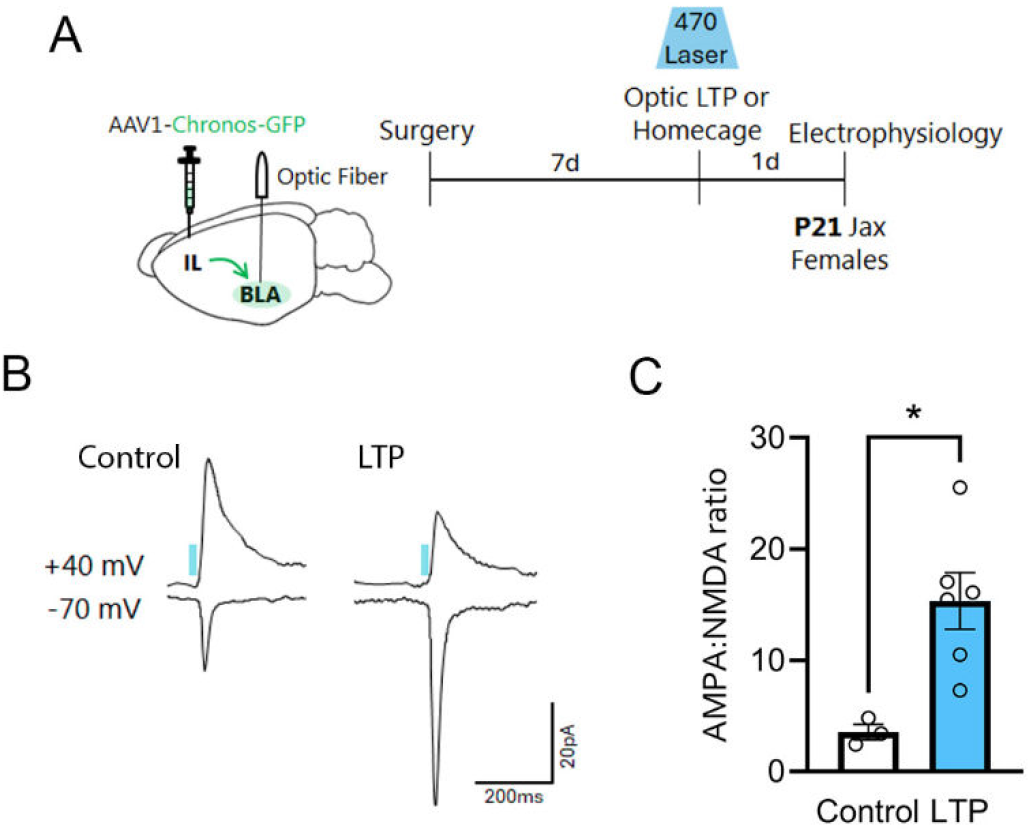
Electrophysiological validation of LTP experiments (Figure 3). **A**. Experimental timeline. Jax female mice were infused with pAAV1-Syn-Chronos-GFP into the IL and implanted with optic fibers over the BLA. At P20, all mice underwent an optic LTP protocol in the homecage, followed by patch clamping of BLA principal neurons the next day (P21). **B**. Representative excitatory postsynaptic current (EPSC) traces resulting from optic stimulation at -70mV or +40mV. For calculation of AMPA/NMDA ratio (**C**), AMPAR-EPSC amplitude was defined as the peak amplitude at -70 mV, when NMDA receptors contribute negligible current. NMDAR-EPSCs were measured at +40 mV at 100ms after LED stimulation, when AMPAR currents have fully decayed and responses are dominated by NMDAR-mediated currents. **C**. mPFC-BLA optic LTP resulted in an increase in AMPA:NMDA ratio in mPFC-BLA synapses (unpaired t test, t_7=_3.132, p= 0.016). (n=9 cells, 5 mice). *p<0.05.

**Table S1.**
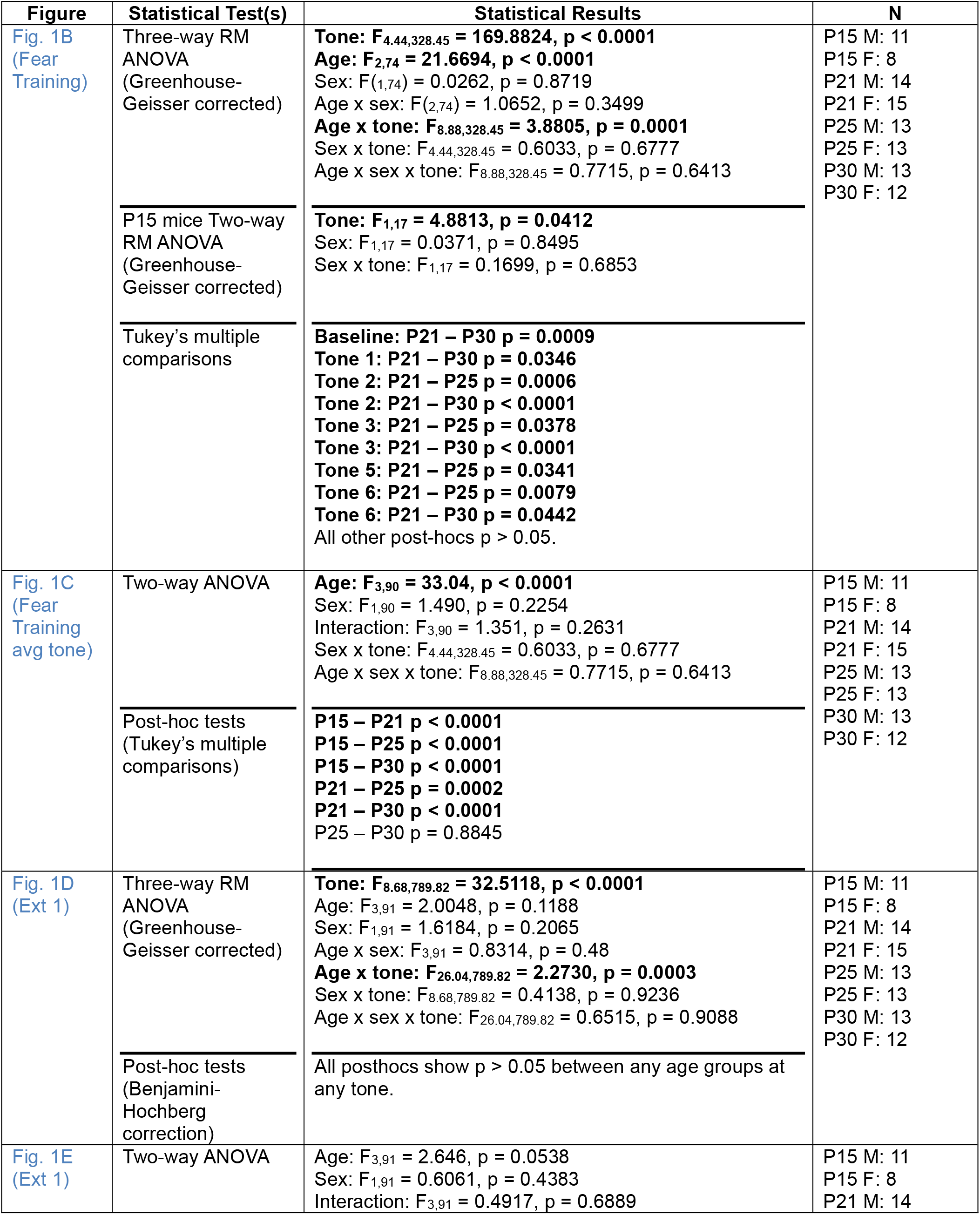
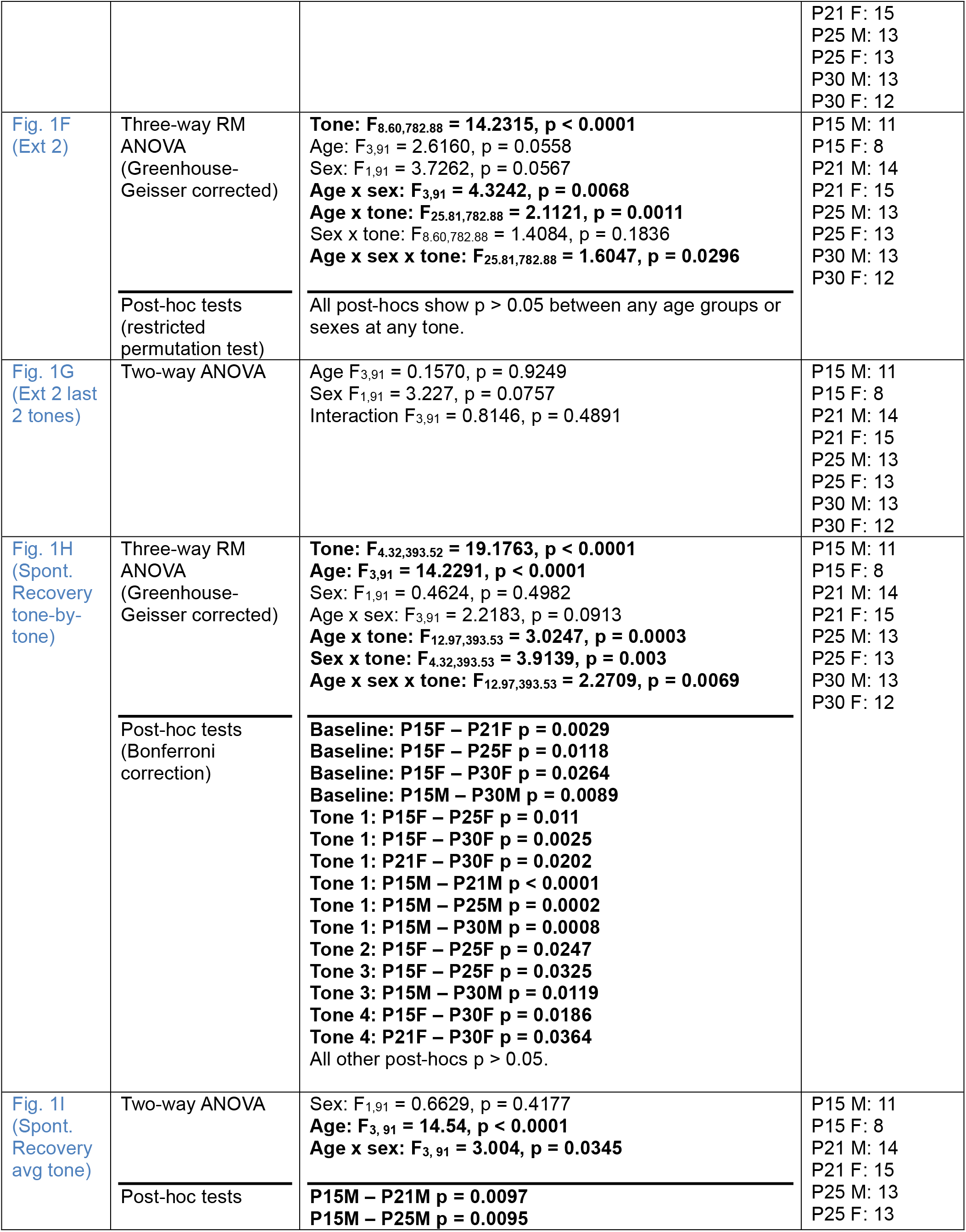
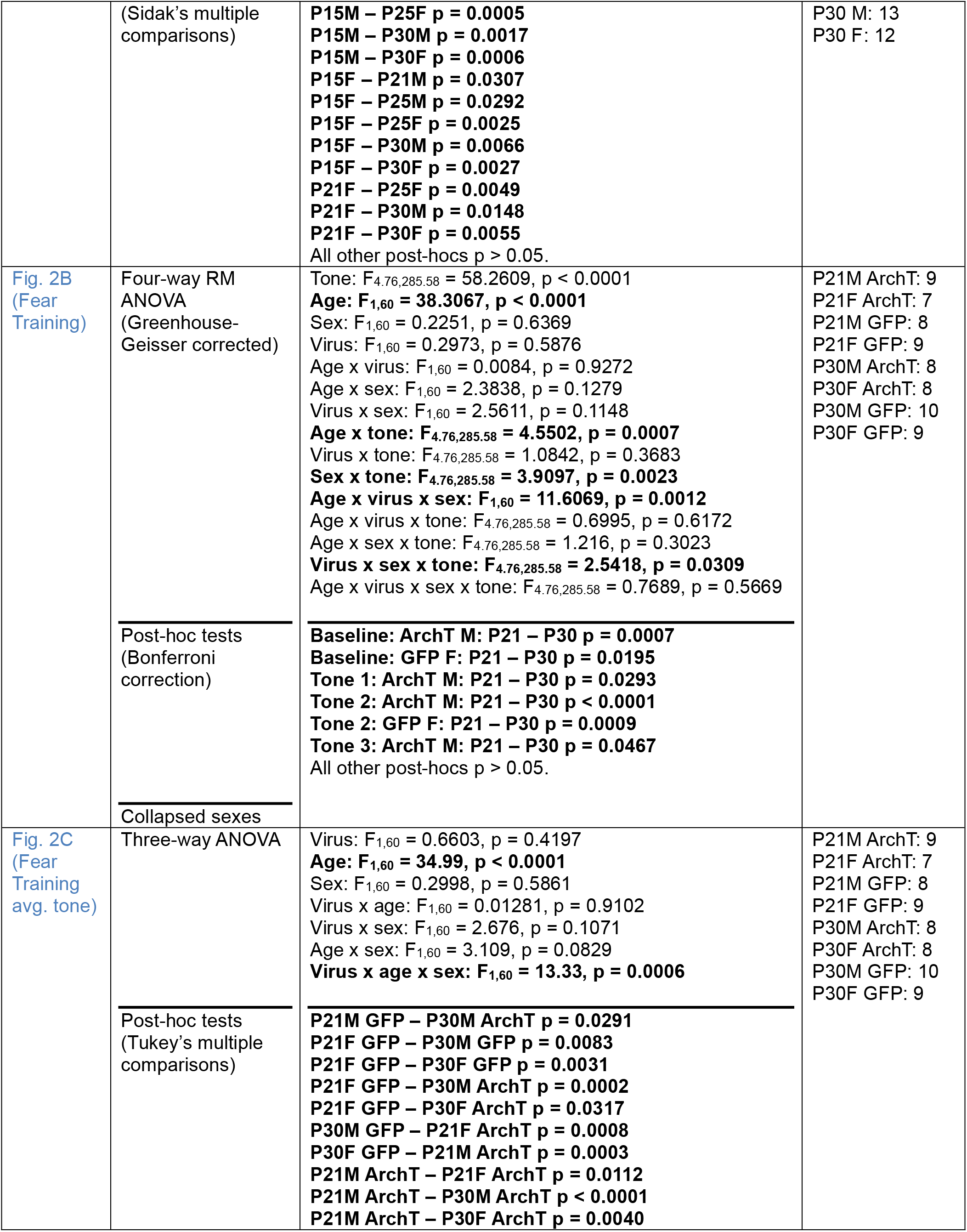
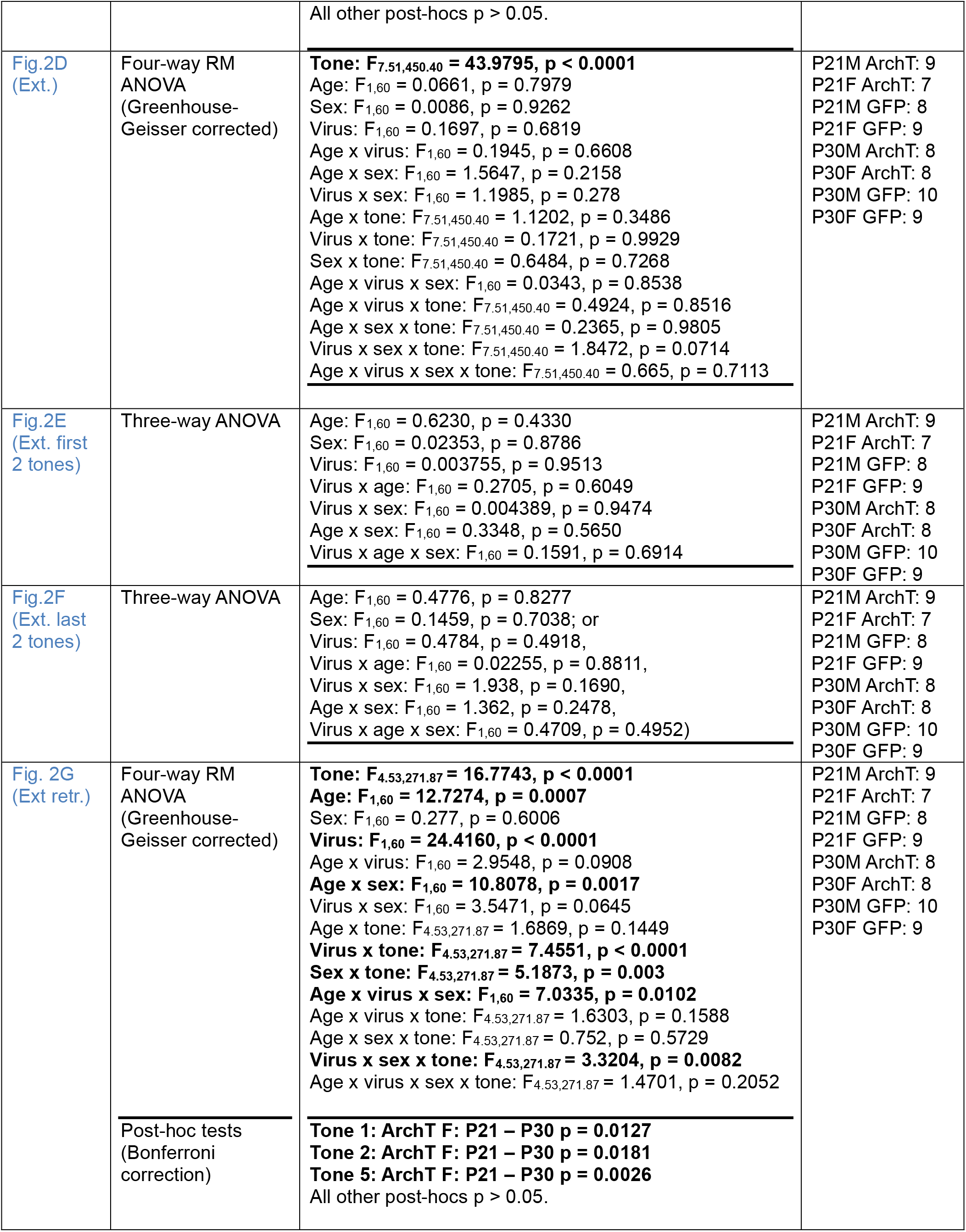
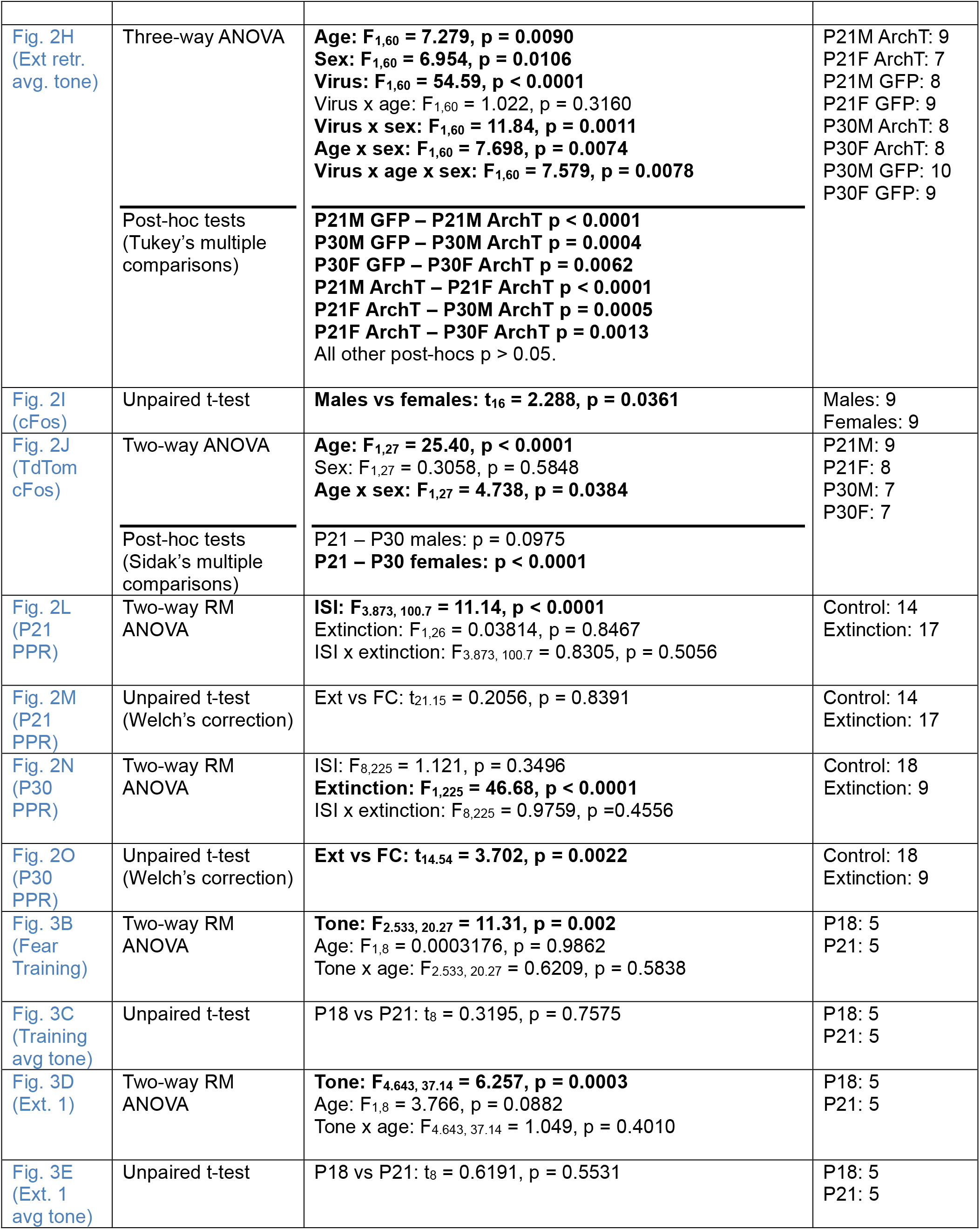
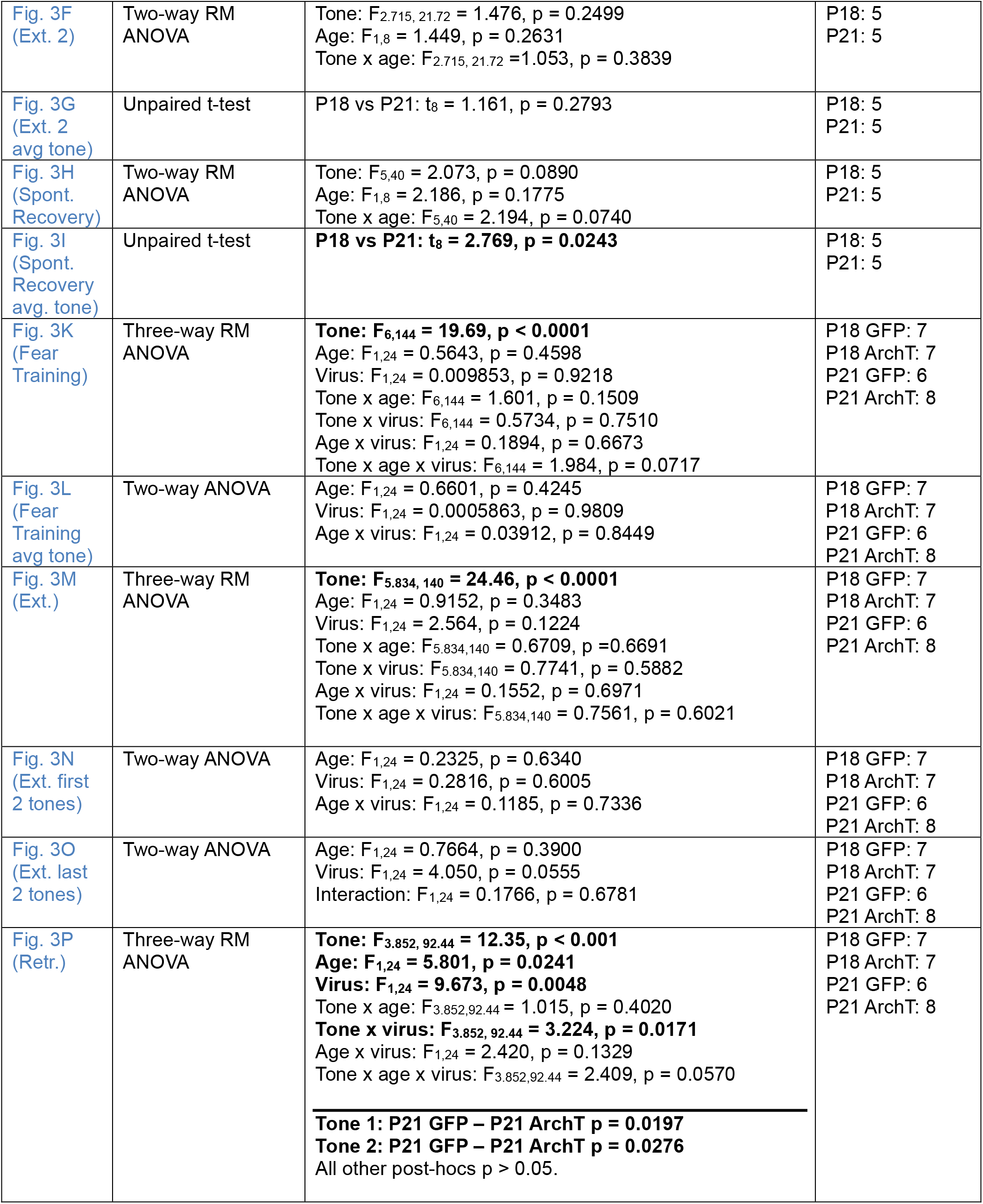
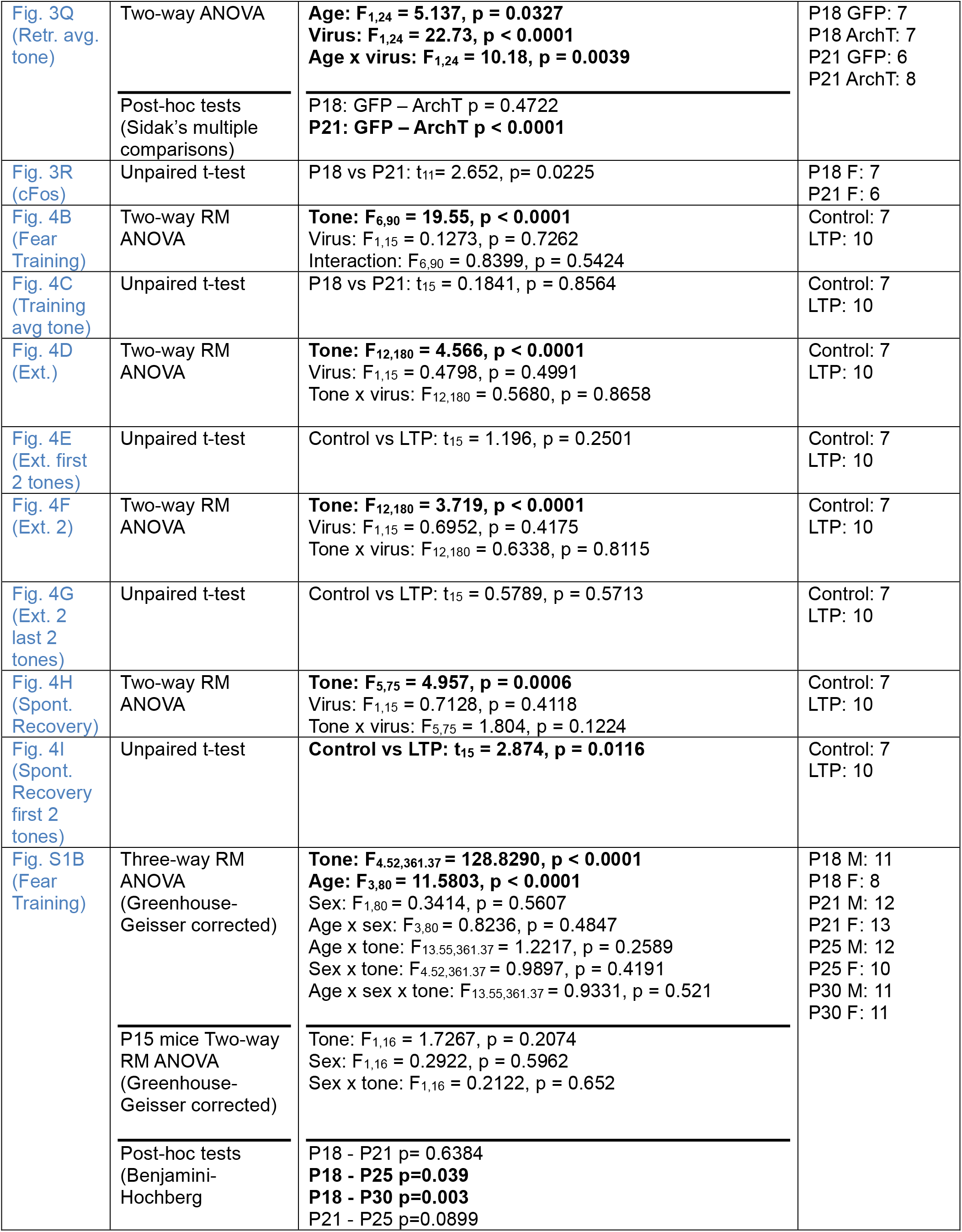
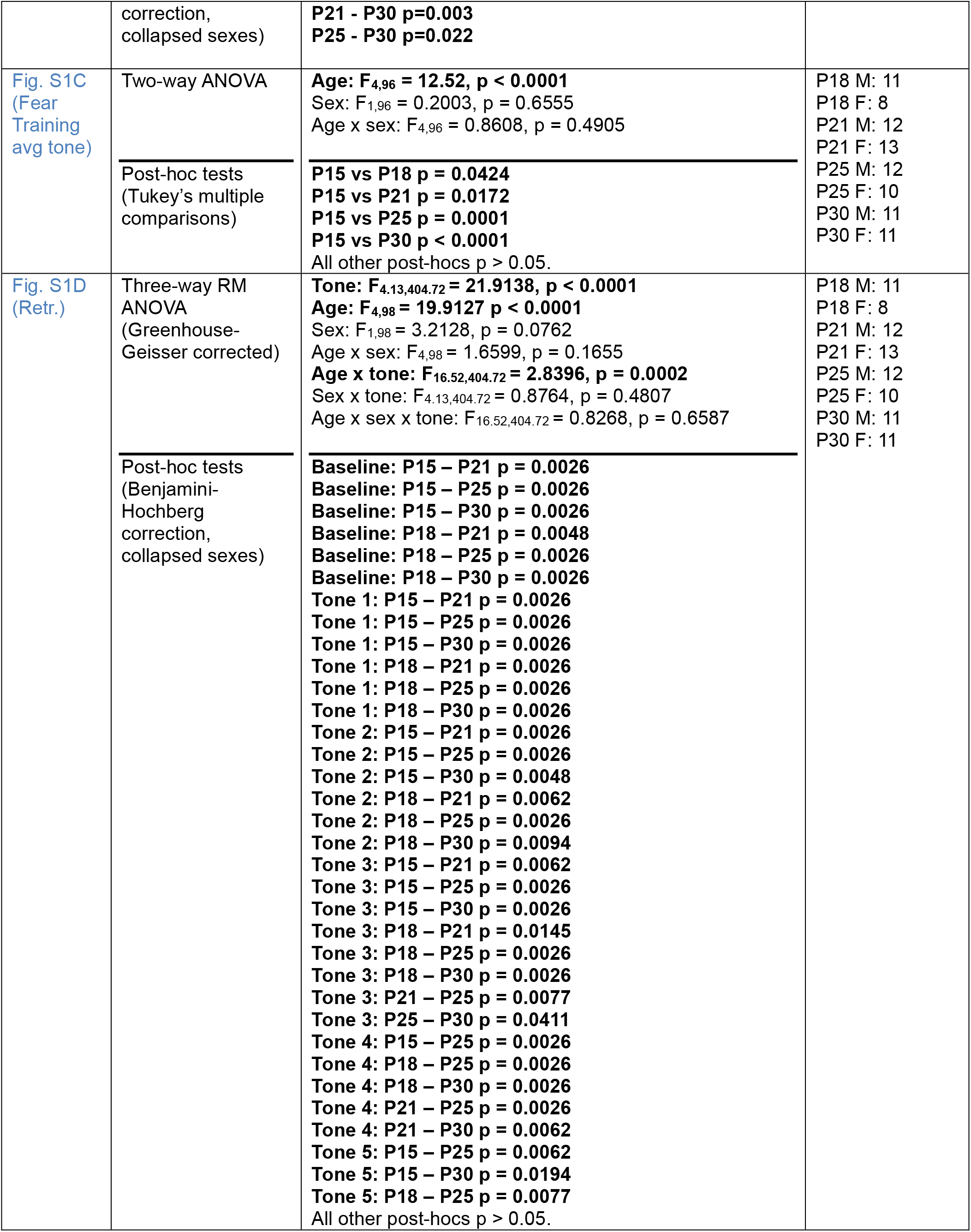
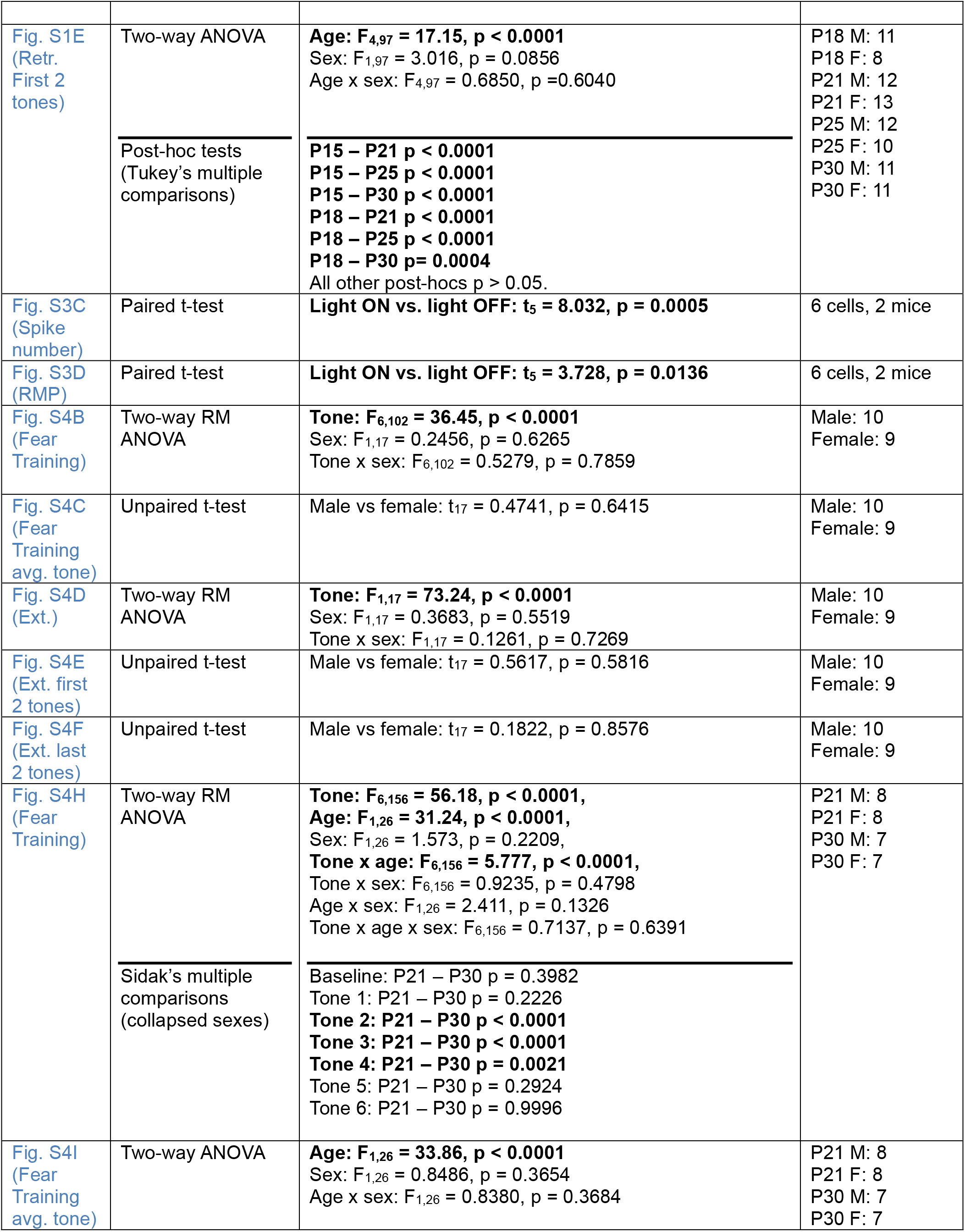
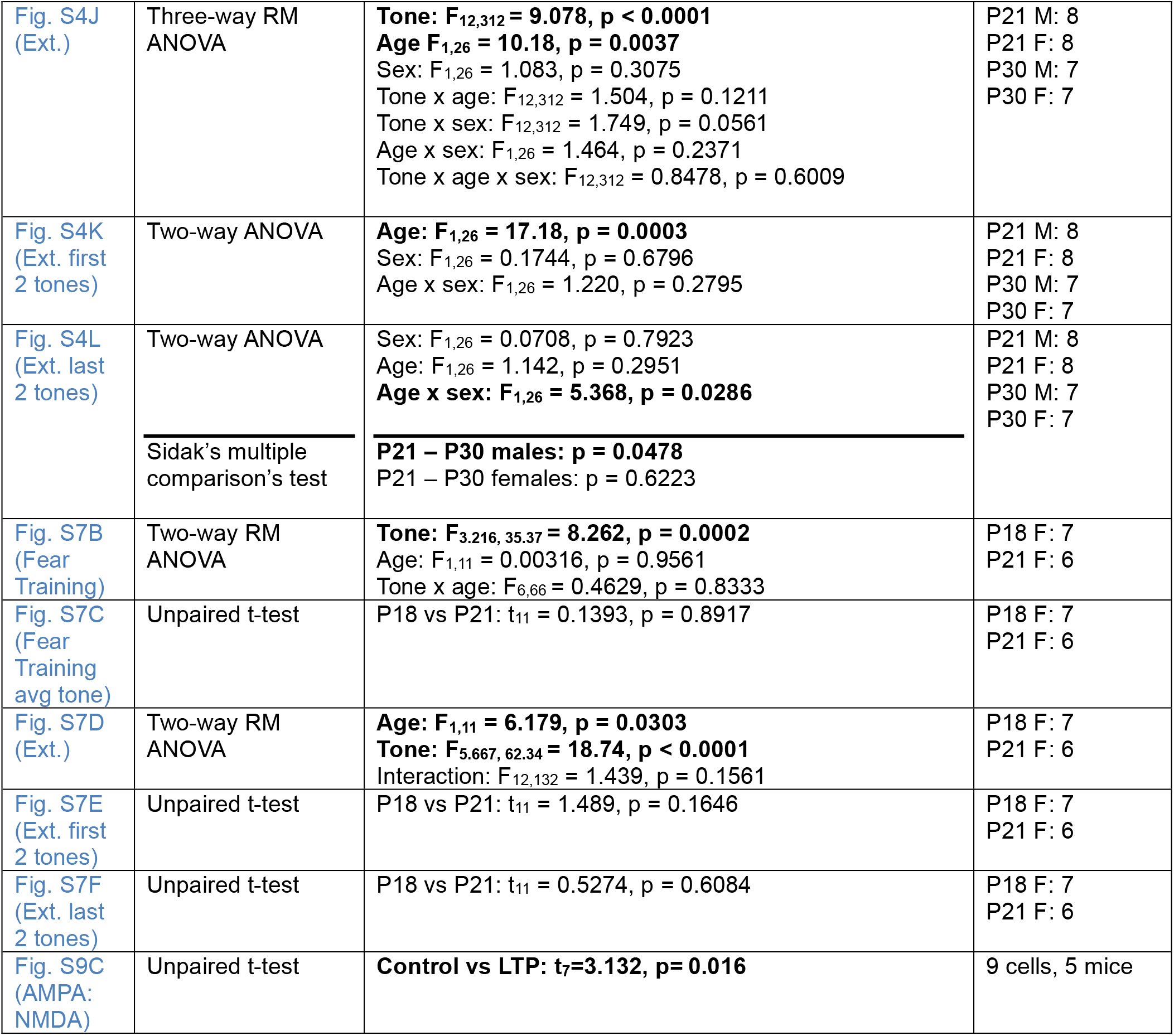
Detailed statistical analysis. Bolded results are significant at α = 0.05.

| Figure | Statistical Test(s) | Statistical Results | N |
|---|---|---|---|
| Fig. 1B (Fear Training) | Three-way RM ANOVA (Greenhouse-Geisser corrected)<br>P15 mice Two-way RM ANOVA (Greenhouse-Geisser corrected)<br>Tukey’s multiple comparisons | **Tone: F<sub>4.44,328.45</sub> = 169.8824, p < 0.0001**<br>**Age: F<sub>2,74</sub> = 21.6694, p < 0.0001**<br>Sex: F(<sub>1,74</sub>) = 0.0262, p = 0.8719<br>Age x sex: F(<sub>2,74</sub>) = 1.0652, p = 0.3499<br>**Age x tone: F<sub>8.88,328.45</sub> = 3.8805, p = 0.0001**<br>Sex x tone: F<sub>4.44,328.45</sub> = 0.6033, p = 0.6777<br>Age x sex x tone: F<sub>8.88,328.45</sub> = 0.7715, p = 0.6413<br>**Tone: F<sub>1,17</sub> = 4.8813, p = 0.0412**<br>Sex: F<sub>1,17</sub> = 0.0371, p = 0.8495<br>Sex x tone: F<sub>1,17</sub> = 0.1699, p = 0.6853<br>**Baseline: P21 – P30 p = 0.0009**<br>**Tone 1: P21 – P30 p = 0.0346**<br>**Tone 2: P21 – P25 p = 0.0006**<br>**Tone 2: P21 – P30 p < 0.0001**<br>**Tone 3: P21 – P25 p = 0.0378**<br>**Tone 3: P21 – P30 p < 0.0001**<br>**Tone 5: P21 – P25 p = 0.0341**<br>**Tone 6: P21 – P25 p = 0.0079**<br>**Tone 6: P21 – P30 p = 0.0442**<br>All other post-hocs p > 0.05. | P15 M: 11<br>P15 F: 8<br>P21 M: 14<br>P21 F: 15<br>P25 M: 13<br>P25 F: 13<br>P30 M: 13<br>P30 F: 12 |
| Fig. 1C (Fear Training avg tone) | Two-way ANOVA<br>Post-hoc tests (Tukey’s multiple comparisons) | **Age: F<sub>3,90</sub> = 33.04, p < 0.0001**<br>Sex: F<sub>1,90</sub> = 1.490, p = 0.2254<br>Interaction: F<sub>3,90</sub> = 1.351, p = 0.2631<br>Sex x tone: F<sub>4.44,328.45</sub> = 0.6033, p = 0.6777<br>Age x sex x tone: F<sub>8.88,328.45</sub> = 0.7715, p = 0.6413<br>**P15 – P21 p < 0.0001**<br>**P15 – P25 p < 0.0001**<br>**P15 – P30 p < 0.0001**<br>**P21 – P25 p = 0.0002**<br>**P21 – P30 p < 0.0001**<br>P25 – P30 p = 0.8845 | P15 M: 11<br>P15 F: 8<br>P21 M: 14<br>P21 F: 15<br>P25 M: 13<br>P25 F: 13<br>P30 M: 13<br>P30 F: 12 |
| Fig. 1D (Ext 1) | Three-way RM ANOVA (Greenhouse-Geisser corrected)<br>Post-hoc tests (Benjamini-Hochberg correction) | **Tone: F<sub>8.68,789.82</sub> = 32.5118, p < 0.0001**<br>Age: F<sub>3,91</sub> = 2.0048, p = 0.1188<br>Sex: F<sub>1,91</sub> = 1.6184, p = 0.2065<br>Age x sex: F<sub>3,91</sub> = 0.8314, p = 0.48<br>**Age x tone: F<sub>26.04,789.82</sub> = 2.2730, p = 0.0003**<br>Sex x tone: F<sub>8.68,789.82</sub> = 0.4138, p = 0.9236<br>Age x sex x tone: F<sub>26.04,789.82</sub> = 0.6515, p = 0.9088<br>All posthocs show p > 0.05 between any age groups at any tone. | P15 M: 11<br>P15 F: 8<br>P21 M: 14<br>P21 F: 15<br>P25 M: 13<br>P25 F: 13<br>P30 M: 13<br>P30 F: 12 |
| Fig. 1E (Ext 1) | Two-way ANOVA | Age: F<sub>3,91</sub> = 2.646, p = 0.0538<br>Sex: F<sub>1,91</sub> = 0.6061, p = 0.4383<br>Interaction: F<sub>3,91</sub> = 0.4917, p = 0.6889 | P15 M: 11<br>P15 F: 8<br>P21 M: 14<br>P21 F: 15<br>P25 M: 13<br>P25 F: 13<br>P30 M: 13<br>P30 F: 12 |
| Fig. 1F (Ext 2) | Three-way RM ANOVA (Greenhouse-Geisser corrected)<br>Post-hoc tests (restricted permutation test) | **Tone: F<sub>8.60,782.88</sub> = 14.2315, p < 0.0001**<br>Age: F<sub>3,91</sub> = 2.6160, p = 0.0558<br>Sex: F<sub>1,91</sub> = 3.7262, p = 0.0567<br>**Age x sex: F<sub>3,91</sub> = 4.3242, p = 0.0068**<br>**Age x tone: F<sub>25.81,782.88</sub> = 2.1121, p = 0.0011**<br>Sex x tone: F<sub>8.60,782.88</sub> = 1.4084, p = 0.1836<br>**Age x sex x tone: F<sub>25.81,782.88</sub> = 1.6047, p = 0.0296**<br>All post-hocs show p > 0.05 between any age groups or sexes at any tone. | P15 M: 11<br>P15 F: 8<br>P21 M: 14<br>P21 F: 15<br>P25 M: 13<br>P25 F: 13<br>P30 M: 13<br>P30 F: 12 |
| Fig. 1G (Ext 2 last 2 tones) | Two-way ANOVA | Age F<sub>3,91</sub> = 0.1570, p = 0.9249<br>Sex F<sub>1,91</sub> = 3.227, p = 0.0757<br>Interaction F<sub>3,91</sub> = 0.8146, p = 0.4891 | P15 M: 11<br>P15 F: 8<br>P21 M: 14<br>P21 F: 15<br>P25 M: 13<br>P25 F: 13<br>P30 M: 13<br>P30 F: 12 |
| Fig. 1H (Spont. Recovery tone-by-tone) | Three-way RM ANOVA (Greenhouse-Geisser corrected)<br>Post-hoc tests (Bonferroni correction) | **Tone: F<sub>4.32,393.52</sub> = 19.1763, p < 0.0001**<br>**Age: F<sub>3,91</sub> = 14.2291, p < 0.0001**<br>Sex: F<sub>1,91</sub> = 0.4624, p = 0.4982<br>Age x sex: F<sub>3,91</sub> = 2.2183, p = 0.0913<br>**Age x tone: F<sub>12.97,393.53</sub> = 3.0247, p = 0.0003**<br>**Sex x tone: F<sub>4.32,393.53</sub> = 3.9139, p = 0.003**<br>**Age x sex x tone: F<sub>12.97,393.53</sub> = 2.2709, p = 0.0069**<br>**Baseline: P15F – P21F p = 0.0029**<br>**Baseline: P15F – P25F p = 0.0118**<br>**Baseline: P15F – P30F p = 0.0264**<br>**Baseline: P15M – P30M p = 0.0089**<br>**Tone 1: P15F – P25F p = 0.011**<br>**Tone 1: P15F – P30F p = 0.0025**<br>**Tone 1: P21F – P30F p = 0.0202**<br>**Tone 1: P15M – P21M p < 0.0001**<br>**Tone 1: P15M – P25M p = 0.0002**<br>**Tone 1: P15M – P30M p = 0.0008**<br>**Tone 2: P15F – P25F p = 0.0247**<br>**Tone 3: P15F – P25F p = 0.0325**<br>**Tone 3: P15M – P30M p = 0.0119**<br>**Tone 4: P15F – P30F p = 0.0186**<br>**Tone 4: P21F – P30F p = 0.0364**<br>All other post-hocs p > 0.05. | P15 M: 11<br>P15 F: 8<br>P21 M: 14<br>P21 F: 15<br>P25 M: 13<br>P25 F: 13<br>P30 M: 13<br>P30 F: 12 |
| Fig. 1I (Spont. Recovery avg tone) | Two-way ANOVA<br>Post-hoc tests (Sidak’s multiple comparisons) | Sex: F<sub>1,91</sub> = 0.6629, p = 0.4177<br>**Age: F<sub>3, 91</sub> = 14.54, p < 0.0001**<br>**Age x sex: F<sub>3, 91</sub> = 3.004, p = 0.0345**<br>**P15M – P21M p = 0.0097**<br>**P15M – P25M p = 0.0095**<br>**P15M – P25F p = 0.0005**<br>**P15M – P30M p = 0.0017**<br>**P15M – P30F p = 0.0006**<br>**P15F – P21M p = 0.0307**<br>**P15F – P25M p = 0.0292**<br>**P15F – P25F p = 0.0025**<br>**P15F – P30M p = 0.0066**<br>**P15F – P30F p = 0.0027**<br>**P21F – P25F p = 0.0049**<br>**P21F – P30M p = 0.0148**<br>**P21F – P30F p = 0.0055**<br>All other post-hocs p > 0.05. | P15 M: 11<br>P15 F: 8<br>P21 M: 14<br>P21 F: 15<br>P25 M: 13<br>P25 F: 13<br>P30 M: 13<br>P30 F: 12 |
| Fig. 2B (Fear Training) | Four-way RM ANOVA (Greenhouse-Geisser corrected)<br>Post-hoc tests (Bonferroni correction)<br>Collapsed sexes | Tone: F<sub>4.76,285.58</sub> = 58.2609, p < 0.0001<br>**Age: F<sub>1,60</sub> = 38.3067, p < 0.0001**<br>Sex: F<sub>1,60</sub> = 0.2251, p = 0.6369<br>Virus: F<sub>1,60</sub> = 0.2973, p = 0.5876<br>Age x virus: F<sub>1,60</sub> = 0.0084, p = 0.9272<br>Age x sex: F<sub>1,60</sub> = 2.3838, p = 0.1279<br>Virus x sex: F<sub>1,60</sub> = 2.5611, p = 0.1148<br>**Age x tone: F<sub>4.76,285.58</sub> = 4.5502, p = 0.0007**<br>Virus x tone: F<sub>4.76,285.58</sub> = 1.0842, p = 0.3683<br>**Sex x tone: F<sub>4.76,285.58</sub> = 3.9097, p = 0.0023**<br>**Age x virus x sex: F<sub>1,60</sub> = 11.6069, p = 0.0012**<br>Age x virus x tone: F<sub>4.76,285.58</sub> = 0.6995, p = 0.6172<br>Age x sex x tone: F<sub>4.76,285.58</sub> = 1.216, p = 0.3023<br>**Virus x sex x tone: F<sub>4.76,285.58</sub> = 2.5418, p = 0.0309**<br>Age x virus x sex x tone: F<sub>4.76,285.58</sub> = 0.7689, p = 0.5669<br>**Baseline: ArchT M: P21 – P30 p = 0.0007**<br>**Baseline: GFP F: P21 – P30 p = 0.0195**<br>**Tone 1: ArchT M: P21 – P30 p = 0.0293**<br>**Tone 2: ArchT M: P21 – P30 p < 0.0001**<br>**Tone 2: GFP F: P21 – P30 p = 0.0009**<br>**Tone 3: ArchT M: P21 – P30 p = 0.0467**<br>All other post-hocs p > 0.05. | P21M ArchT: 9<br>P21F ArchT: 7<br>P21M GFP: 8<br>P21F GFP: 9<br>P30M ArchT: 8<br>P30F ArchT: 8<br>P30M GFP: 10<br>P30F GFP: 9 |
| Fig. 2C (Fear Training avg. tone) | Three-way ANOVA<br>Post-hoc tests (Tukey’s multiple comparisons) | Virus: F<sub>1,60</sub> = 0.6603, p = 0.4197<br>**Age: F<sub>1,60</sub> = 34.99, p < 0.0001**<br>Sex: F<sub>1,60</sub> = 0.2998, p = 0.5861<br>Virus x age: F<sub>1,60</sub> = 0.01281, p = 0.9102<br>Virus x sex: F<sub>1,60</sub> = 2.676, p = 0.1071<br>Age x sex: F<sub>1,60</sub> = 3.109, p = 0.0829<br>**Virus x age sex: F<sub>1,60</sub> = 13.33, p = 0.0006**<br>**P21M GFP – P30M ArchT p = 0.0291**<br>**P21F GFP – P30M GFP p = 0.0083**<br>**P21F GFP – P30F GFP p = 0.0031**<br>**P21F GFP – P30M ArchT p = 0.0002**<br>**P21F GFP – P30F ArchT p = 0.0317**<br>**P30M GFP – P21F ArchT p = 0.0008**<br>**P30F GFP – P21M ArchT p = 0.0003**<br>**P21M ArchT – P21F ArchT p = 0.0112**<br>**P21M ArchT – P30M ArchT p < 0.0001**<br>**P21M ArchT – P30F ArchT p = 0.0040**<br>All other post-hocs p > 0.05. | P21M ArchT: 9<br>P21F ArchT: 7<br>P21M GFP: 8<br>P21F GFP: 9<br>P30M ArchT: 8<br>P30F ArchT: 8<br>P30M GFP: 10<br>P30F GFP: 9 |
| Fig.2D (Ext.) | Four-way RM ANOVA (Greenhouse-Geisser corrected) | **Tone: F<sub>7.51,450.40</sub> = 43.9795, p < 0.0001**<br>Age: F<sub>1,60</sub> = 0.0661, p = 0.7979<br>Sex: F<sub>1,60</sub> = 0.0086, p = 0.9262<br>Virus: F<sub>1,60</sub> = 0.1697, p = 0.6819<br>Age x virus: F<sub>1,60</sub> = 0.1945, p = 0.6608<br>Age x sex: F<sub>1,60</sub> = 1.5647, p = 0.2158<br>Virus x sex: F<sub>1,60</sub> = 1.1985, p = 0.278<br>Age x tone: F<sub>7.51,450.40</sub> = 1.1202, p = 0.3486<br>Virus x tone: F<sub>7.51,450.40</sub> = 0.1721, p = 0.9929<br>Sex x tone: F<sub>7.51,450.40</sub> = 0.6484, p = 0.7268<br>Age x virus x sex: F<sub>1,60</sub> = 0.0343, p = 0.8538<br>Age x virus x tone: F<sub>7.51,450.40</sub> = 0.4924, p = 0.8516<br>Age x sex x tone: F<sub>7.51,450.40</sub> = 0.2365, p = 0.9805<br>Virus x sex x tone: F<sub>7.51,450.40</sub> = 1.8472, p = 0.0714<br>Age x virus x sex x tone: F<sub>7.51,450.40</sub> = 0.665, p = 0.7113 | P21M ArchT: 9<br>P21F ArchT: 7<br>P21M GFP: 8<br>P21F GFP: 9<br>P30M ArchT: 8<br>P30F ArchT: 8<br>P30M GFP: 10<br>P30F GFP: 9 |
| Fig.2E (Ext. first 2 tones) | Three-way ANOVA | Age: F<sub>1,60</sub> = 0.6230, p = 0.4330<br>Sex: F<sub>1,60</sub> = 0.02353, p = 0.8786<br>Virus: F<sub>1,60</sub> = 0.003755, p = 0.9513<br>Virus x age: F<sub>1,60</sub> = 0.2705, p = 0.6049<br>Virus x sex: F<sub>1,60</sub> = 0.004389, p = 0.9474<br>Age x sex: F<sub>1,60</sub> = 0.3348, p = 0.5650<br>Virus x age x sex: F<sub>1,60</sub> = 0.1591, p = 0.6914 | P21M ArchT: 9<br>P21F ArchT: 7<br>P21M GFP: 8<br>P21F GFP: 9<br>P30M ArchT: 8<br>P30F ArchT: 8<br>P30M GFP: 10<br>P30F GFP: 9 |
| Fig.2F (Ext. last 2 tones) | Three-way ANOVA | Age: F<sub>1,60</sub> = 0.4776, p = 0.8277<br>Sex: F<sub>1,60</sub> = 0.1459, p = 0.7038; or<br>Virus: F<sub>1,60</sub> = 0.4784, p = 0.4918,<br>Virus x age: F<sub>1,60</sub> = 0.02255, p = 0.8811,<br>Virus x sex: F<sub>1,60</sub> = 1.938, p = 0.1690,<br>Age x sex: F<sub>1,60</sub> = 1.362, p = 0.2478,<br>Virus x age x sex: F<sub>1,60</sub> = 0.4709, p = 0.4952) | P21M ArchT: 9<br>P21F ArchT: 7<br>P21M GFP: 8<br>P21F GFP: 9<br>P30M ArchT: 8<br>P30F ArchT: 8<br>P30M GFP: 10<br>P30F GFP: 9 |
| Fig. 2G (Ext retr.) | Four-way RM ANOVA (Greenhouse-Geisser corrected)<br>Post-hoc tests (Bonferroni correction) | **Tone: F<sub>4.53,271.87</sub> = 16.7743, p < 0.0001**<br>**Age: F<sub>1,60</sub> = 12.7274, p = 0.0007**<br>Sex: F<sub>1,60</sub> = 0.277, p = 0.6006<br>**Virus: F<sub>1,60</sub> = 24.4160, p < 0.0001**<br>Age x virus: F<sub>1,60</sub> = 2.9548, p = 0.0908<br>**Age x sex: F<sub>1,60</sub> = 10.8078, p = 0.0017**<br>Virus x sex: F<sub>1,60</sub> = 3.5471, p = 0.0645<br>Age x tone: F<sub>4.53,271.87</sub> = 1.6869, p = 0.1449<br>**Virus x tone: F<sub>4.53,271.87</sub> = 7.4551, p < 0.0001**<br>**Sex x tone: F<sub>4.53,271.87</sub> = 5.1873, p = 0.003**<br>**Age x virus x sex: F<sub>1,60</sub> = 7.0335, p = 0.0102**<br>Age x virus x tone: F<sub>4.53,271.87</sub> = 1.6303, p = 0.1588<br>Age x sex x tone: F<sub>4.53,271.87</sub> = 0.752, p = 0.5729<br>**Virus x sex x tone: F<sub>4.53,271.87</sub> = 3.3204, p = 0.0082**<br>Age x virus x sex x tone: F<sub>4.53,271.87</sub> = 1.4701, p = 0.2052<br>**Tone 1: ArchT F: P21 – P30 p = 0.0127**<br>**Tone 2: ArchT F: P21 – P30 p = 0.0181**<br>**Tone 5: ArchT F: P21 – P30 p = 0.0026**<br>All other post-hocs p > 0.05. | P21M ArchT: 9<br>P21F ArchT: 7<br>P21M GFP: 8<br>P21F GFP: 9<br>P30M ArchT: 8<br>P30F ArchT: 8<br>P30M GFP: 10<br>P30F GFP: 9 |
| Fig. 2H (Ext retr. avg. tone) | Three-way ANOVA<br>Post-hoc tests (Tukey’s multiple comparisons) | **Age: F<sub>1,60</sub> = 7.279, p = 0.0090**<br>**Sex: F<sub>1,60</sub> = 6.954, p = 0.0106**<br>**Virus: F<sub>1,60</sub> = 54.59, p < 0.0001**<br>Virus x age: F<sub>1,60</sub> = 1.022, p = 0.3160<br>**Virus x sex: F<sub>1,60</sub> = 11.84, p = 0.0011**<br>**Age x sex: F<sub>1,60</sub> = 7.698, p = 0.0074**<br>**Virus x age x sex: F<sub>1,60</sub> = 7.579, p = 0.0078**<br>**P21M GFP – P21M ArchT p < 0.0001**<br>**P30M GFP – P30M ArchT p = 0.0004**<br>**P30F GFP – P30F ArchT p = 0.0062**<br>**P21M ArchT – P21F ArchT p < 0.0001**<br>**P21F ArchT – P30M ArchT p = 0.0005**<br>**P21F ArchT – P30F ArchT p = 0.0013**<br>All other post-hocs p > 0.05. | P21M ArchT: 9<br>P21F ArchT: 7<br>P21M GFP: 8<br>P21F GFP: 9<br>P30M ArchT: 8<br>P30F ArchT: 8<br>P30M GFP: 10<br>P30F GFP: 9 |
| Fig. 2I (cFos) | Unpaired t-test | **Males vs females: t<sub>16</sub> = 2.288, p = 0.0361** | Males: 9<br>Females: 9 |
| Fig. 2J (TdTom cFos) | Two-way ANOVA<br>Post-hoc tests (Sidak’s multiple comparisons) | **Age: F<sub>1,27</sub> = 25.40, p < 0.0001**<br>Sex: F<sub>1,27</sub> = 0.3058, p = 0.5848<br>**Age x sex: F<sub>1,27</sub> = 4.738, p = 0.0384**<br>P21 – P30 males: p = 0.0975<br>**P21 – P30 females: p < 0.0001** | P21M: 9<br>P21F: 8<br>P30M: 7<br>P30F: 7 |
| Fig. 2L (P21 PPR) | Two-way RM ANOVA | **ISI: F<sub>3.873, 100.7</sub> = 11.14, p < 0.0001**<br>Extinction: F<sub>1,26</sub> = 0.03814, p = 0.8467<br>ISI x extinction: F<sub>3.873, 100.7</sub> = 0.8305, p = 0.5056 | Control: 14<br>Extinction: 17 |
| Fig. 2M (P21 PPR) | Unpaired t-test (Welch’s correction) | Ext vs FC: t<sub>21.15</sub> = 0.2056, p = 0.8391 | Control: 14<br>Extinction: 17 |
| Fig. 2N (P30 PPR) | Two-way RM ANOVA | ISI: F<sub>8,225</sub> = 1.121, p = 0.3496<br>**Extinction: F<sub>1,225</sub> = 46.68, p < 0.0001**<br>ISI x extinction: F<sub>8,225</sub> = 0.9759, p =0.4556 | Control: 18<br>Extinction: 9 |
| Fig. 2O (P30 PPR) | Unpaired t-test (Welch’s correction) | **Ext vs FC: t<sub>14.54</sub> = 3.702, p = 0.0022** | Control: 18<br>Extinction: 9 |
| Fig. 3B (Fear Training) | Two-way RM ANOVA | **Tone: F<sub>2.533, 20.27</sub> = 11.31, p = 0.002**<br>Age: F<sub>1,8</sub> = 0.0003176, p = 0.9862<br>Tone x age: F<sub>2.533, 20.27</sub> = 0.6209, p = 0.5838 | P18: 5<br>P21: 5 |
| Fig. 3C (Training avg tone) | Unpaired t-test | P18 vs P21: t<sub>8</sub> = 0.3195, p = 0.7575 | P18: 5<br>P21: 5 |
| Fig. 3D (Ext. 1) | Two-way RM ANOVA | **Tone: F<sub>4.643, 37.14</sub> = 6.257, p = 0.0003**<br>Age: F<sub>1,8</sub> = 3.766, p = 0.0882<br>Tone x age: F<sub>4.643, 37.14</sub> = 1.049, p = 0.4010 | P18: 5<br>P21: 5 |
| Fig. 3E (Ext. 1 avg tone) | Unpaired t-test | P18 vs P21: t<sub>8</sub> = 0.6191, p = 0.5531 | P18: 5<br>P21: 5 |
| Fig. 3F (Ext. 2) | Two-way RM ANOVA | Tone: F<sub>2.715, 21.72</sub> = 1.476, p = 0.2499<br>Age: F<sub>1,8</sub> = 1.449, p = 0.2631<br>Tone x age: F<sub>2.715, 21.72</sub> =1.053, p = 0.3839 | P18: 5<br>P21: 5 |
| Fig. 3G (Ext. 2 avg tone) | Unpaired t-test | P18 vs P21: t<sub>8</sub> = 1.161, p = 0.2793 | P18: 5<br>P21: 5 |
| Fig. 3H (Spont. Recovery) | Two-way RM ANOVA | Tone: F<sub>5,40</sub> = 2.073, p = 0.0890<br>Age: F<sub>1,8</sub> = 2.186, p = 0.1775<br>Tone x age: F<sub>5,40</sub> = 2.194, p = 0.0740 | P18: 5<br>P21: 5 |
| Fig. 3I (Spont. Recovery avg. tone) | Unpaired t-test | **P18 vs P21: t<sub>8</sub> = 2.769, p = 0.0243** | P18: 5<br>P21: 5 |
| Fig. 3K (Fear Training) | Three-way RM ANOVA | **Tone: F<sub>6,144</sub> = 19.69, p < 0.0001**<br>Age: F<sub>1,24</sub> = 0.5643, p = 0.4598<br>Virus: F<sub>1,24</sub> = 0.009853, p = 0.9218<br>Tone x age: F<sub>6,144</sub> = 1.601, p = 0.1509<br>Tone x virus: F<sub>6,144</sub> = 0.5734, p = 0.7510<br>Age x virus: F<sub>1,24</sub> = 0.1894, p = 0.6673<br>Tone x age x virus: F<sub>6,144</sub> = 1.984, p = 0.0717 | P18 GFP: 7<br>P18 ArchT: 7<br>P21 GFP: 6<br>P21 ArchT: 8 |
| Fig. 3L (Fear Training avg tone) | Two-way ANOVA | Age: F<sub>1,24</sub> = 0.6601, p = 0.4245<br>Virus: F<sub>1,24</sub> = 0.0005863, p = 0.9809<br>Age x virus: F<sub>1,24</sub> = 0.03912, p = 0.8449 | P18 GFP: 7<br>P18 ArchT: 7<br>P21 GFP: 6<br>P21 ArchT: 8 |
| Fig. 3M (Ext.) | Three-way RM ANOVA | **Tone: F<sub>5.834, 140</sub> = 24.46, p < 0.0001**<br>Age: F<sub>1,24</sub> = 0.9152, p = 0.3483<br>Virus: F<sub>1,24</sub> = 2.564, p = 0.1224<br>Tone x age: F<sub>5.834,140</sub> = 0.6709, p =0.6691<br>Tone x virus: F<sub>5.834,140</sub> = 0.7741, p = 0.5882<br>Age x virus: F<sub>1,24</sub> = 0.1552, p = 0.6971<br>Tone x age x virus: F<sub>5.834,140</sub> = 0.7561, p = 0.6021 | P18 GFP: 7<br>P18 ArchT: 7<br>P21 GFP: 6<br>P21 ArchT: 8 |
| Fig. 3N (Ext. first 2 tones) | Two-way ANOVA | Age: F<sub>1,24</sub> = 0.2325, p = 0.6340<br>Virus: F<sub>1,24</sub> = 0.2816, p = 0.6005<br>Age x virus: F<sub>1,24</sub> = 0.1185, p = 0.7336 | P18 GFP: 7<br>P18 ArchT: 7<br>P21 GFP: 6<br>P21 ArchT: 8 |
| Fig. 3O (Ext. last 2 tones) | Two-way ANOVA | Age: F<sub>1,24</sub> = 0.7664, p = 0.3900<br>Virus: F<sub>1,24</sub> = 4.050, p = 0.0555<br>Interaction: F<sub>1,24</sub> = 0.1766, p = 0.6781 | P18 GFP: 7<br>P18 ArchT: 7<br>P21 GFP: 6<br>P21 ArchT: 8 |
| Fig. 3P (Retr.) | Three-way RM ANOVA | **Tone: F<sub>3.852, 92.44</sub> = 12.35, p < 0.001**<br>**Age: F<sub>1,24</sub> = 5.801, p = 0.0241**<br>**Virus: F<sub>1,24</sub> = 9.673, p = 0.0048**<br>Tone x age: F<sub>3.852,92.44</sub> = 1.015, p = 0.4020<br>**Tone x virus: F<sub>3.852, 92.44</sub> = 3.224, p = 0.0171**<br>Age x virus: F<sub>1,24</sub> = 2.420, p = 0.1329<br>Tone x age x virus: F<sub>3.852,92.44</sub> = 2.409, p = 0.0570<br>**Tone 1: P21 GFP – P21 ArchT p = 0.0197**<br>**Tone 2: P21 GFP – P21 ArchT p = 0.0276**<br>All other post-hocs p > 0.05. | P18 GFP: 7<br>P18 ArchT: 7<br>P21 GFP: 6<br>P21 ArchT: 8 |
| Fig. 3Q (Retr. avg. tone) | Two-way ANOVA<br>Post-hoc tests (Sidak’s multiple comparisons) | **Age: F<sub>1,24</sub> = 5.137, p = 0.0327**<br>**Virus: F<sub>1,24</sub> = 22.73, p < 0.0001**<br>**Age x virus: F<sub>1,24</sub> = 10.18, p = 0.0039**<br>P18: GFP – ArchT p = 0.4722<br>**P21: GFP – ArchT p < 0.0001** | P18 GFP: 7<br>P18 ArchT: 7<br>P21 GFP: 6<br>P21 ArchT: 8 |
| Fig. 3R (cFos) | Unpaired t-test | P18 vs P21: t<sub>11</sub>= 2.652, p= 0.0225 | P18 F: 7<br>P21 F: 6 |
| Fig. 4B (Fear Training) | Two-way RM ANOVA | **Tone: F<sub>6,90</sub> = 19.55, p < 0.0001**<br>Virus: F<sub>1,15</sub> = 0.1273, p = 0.7262<br>Interaction: F<sub>6,90</sub> = 0.8399, p = 0.5424 | Control: 7<br>LTP: 10 |
| Fig. 4C (Training avg tone) | Unpaired t-test | P18 vs P21: t<sub>15</sub> = 0.1841, p = 0.8564 | Control: 7<br>LTP: 10 |
| Fig. 4D (Ext.) | Two-way RM ANOVA | **Tone: F<sub>12,180</sub> = 4.566, p < 0.0001**<br>Virus: F<sub>1,15</sub> = 0.4798, p = 0.4991<br>Tone x virus: F<sub>12,180</sub> = 0.5680, p = 0.8658 | Control: 7<br>LTP: 10 |
| Fig. 4E (Ext. first 2 tones) | Unpaired t-test | Control vs LTP: t<sub>15</sub> = 1.196, p = 0.2501 | Control: 7<br>LTP: 10 |
| Fig. 4F (Ext. 2) | Two-way RM ANOVA | **Tone: F<sub>12,180</sub> = 3.719, p < 0.0001**<br>Virus: F<sub>1,15</sub> = 0.6952, p = 0.4175<br>Tone x virus: F<sub>12,180</sub> = 0.6338, p = 0.8115 | Control: 7<br>LTP: 10 |
| Fig. 4G (Ext. 2 last 2 tones) | Unpaired t-test | Control vs LTP: t<sub>15</sub> = 0.5789, p = 0.5713 | Control: 7<br>LTP: 10 |
| Fig. 4H (Spont. Recovery) | Two-way RM ANOVA | **Tone: F<sub>5,75</sub> = 4.957, p = 0.0006**<br>Virus: F<sub>1,15</sub> = 0.7128, p = 0.4118<br>Tone x virus: F<sub>5,75</sub> = 1.804, p = 0.1224 | Control: 7<br>LTP: 10 |
| Fig. 4I (Spont. Recovery first 2 tones) | Unpaired t-test | **Control vs LTP: t<sub>15</sub> = 2.874, p = 0.0116** | Control: 7<br>LTP: 10 |
| Fig. S1B (Fear Training) | Three-way RM ANOVA (Greenhouse-Geisser corrected)<br>P15 mice Two-way RM ANOVA (Greenhouse-Geisser corrected)<br>Post-hoc tests (Benjamini-Hochberg correction, collapsed sexes) | **Tone: F<sub>4.52,361.37</sub> = 128.8290, p < 0.0001**<br>**Age: F<sub>3,80</sub> = 11.5803, p < 0.0001**<br>Sex: F<sub>1,80</sub> = 0.3414, p = 0.5607<br>Age x sex: F<sub>3,80</sub> = 0.8236, p = 0.4847<br>Age x tone: F<sub>13.55,361.37</sub> = 1.2217, p = 0.2589<br>Sex x tone: F<sub>4.52,361.37</sub> = 0.9897, p = 0.4191<br>Age x sex x tone: F<sub>13.55,361.37</sub> = 0.9331, p = 0.521<br>Tone: F<sub>1,16</sub> = 1.7267, p = 0.2074<br>Sex: F<sub>1,16</sub> = 0.2922, p = 0.5962<br>Sex x tone: F<sub>1,16</sub> = 0.2122, p = 0.652<br>P18 - P21 p= 0.6384<br>**P18 - P25 p=0.039**<br>**P18 - P30 p=0.003**<br>P21 - P25 p=0.0899<br>**P21 - P30 p=0.003**<br>**P25 - P30 p=0.022** | P18 M: 11<br>P18 F: 8<br>P21 M: 12<br>P21 F: 13<br>P25 M: 12<br>P25 F: 10<br>P30 M: 11<br>P30 F: 11 |
| Fig. S1C (Fear Training avg tone) | Two-way ANOVA<br>Post-hoc tests (Tukey’s multiple comparisons) | **Age: F<sub>4,96</sub> = 12.52, p < 0.0001**<br>Sex: F<sub>1,96</sub> = 0.2003, p = 0.6555<br>Age x sex: F<sub>4,96</sub> = 0.8608, p = 0.4905<br>**P15 vs P18 p = 0.0424**<br>**P15 vs P21 p = 0.0172**<br>**P15 vs P25 p = 0.0001**<br>**P15 vs P30 p < 0.0001**<br>All other post-hocs p > 0.05. | P18 M: 11<br>P18 F: 8<br>P21 M: 12<br>P21 F: 13<br>P25 M: 12<br>P25 F: 10<br>P30 M: 11<br>P30 F: 11 |
| Fig. S1D (Retr.) | Three-way RM ANOVA (Greenhouse-Geisser corrected)<br>Post-hoc tests (Benjamini-Hochberg correction, collapsed sexes) | **Tone: F<sub>4.13,404.72</sub> = 21.9138, p < 0.0001**<br>**Age: F<sub>4,98</sub> = 19.9127 p < 0.0001**<br>Sex: F<sub>1,98</sub> = 3.2128, p = 0.0762<br>Age x sex: F<sub>4,98</sub> = 1.6599, p = 0.1655<br>**Age x tone: F<sub>16.52,404.72</sub> = 2.8396, p = 0.0002**<br>Sex x tone: F<sub>4.13,404.72</sub> = 0.8764, p = 0.4807<br>Age x sex x tone: F<sub>16.52,404.72</sub> = 0.8268, p = 0.6587<br>**Baseline: P15 – P21 p = 0.0026**<br>**Baseline: P15 – P25 p = 0.0026**<br>**Baseline: P15 – P30 p = 0.0026**<br>**Baseline: P18 – P21 p = 0.0048**<br>**Baseline: P18 – P25 p = 0.0026**<br>**Baseline: P18 – P30 p = 0.0026**<br>**Tone 1: P15 – P21 p = 0.0026**<br>**Tone 1: P15 – P25 p = 0.0026**<br>**Tone 1: P15 – P30 p = 0.0026**<br>**Tone 1: P18 – P21 p = 0.0026**<br>**Tone 1: P18 – P25 p = 0.0026**<br>**Tone 1: P18 – P30 p = 0.0026**<br>**Tone 2: P15 – P21 p = 0.0026**<br>**Tone 2: P15 – P25 p = 0.0026**<br>**Tone 2: P15 – P30 p = 0.0048**<br>**Tone 2: P18 – P21 p = 0.0062**<br>**Tone 2: P18 – P25 p = 0.0026**<br>**Tone 2: P18 – P30 p = 0.0094**<br>**Tone 3: P15 – P21 p = 0.0062**<br>**Tone 3: P15 – P25 p = 0.0026**<br>**Tone 3: P15 – P30 p = 0.0026**<br>**Tone 3: P18 – P21 p = 0.0145**<br>**Tone 3: P18 – P25 p = 0.0026**<br>**Tone 3: P18 – P30 p = 0.0026**<br>**Tone 3: P21 – P25 p = 0.0077**<br>**Tone 3: P25 – P30 p = 0.0411**<br>**Tone 4: P15 – P25 p = 0.0026**<br>**Tone 4: P18 – P25 p = 0.0026**<br>**Tone 4: P18 – P30 p = 0.0026**<br>**Tone 4: P21 – P25 p = 0.0026**<br>**Tone 4: P21 – P30 p = 0.0062**<br>**Tone 5: P15 – P25 p = 0.0062**<br>**Tone 5: P15 – P30 p = 0.0194**<br>**Tone 5: P18 – P25 p = 0.0077**<br>All other post-hocs p > 0.05. | P18 M: 11<br>P18 F: 8<br>P21 M: 12<br>P21 F: 13<br>P25 M: 12<br>P25 F: 10<br>P30 M: 11<br>P30 F: 11 |
| Fig. S1E (Retr. First 2 tones) | Two-way ANOVA<br>Post-hoc tests (Tukey’s multiple comparisons) | **Age: F<sub>4,97</sub> = 17.15, p < 0.0001**<br>Sex: F<sub>1,97</sub> = 3.016, p = 0.0856<br>Age x sex: F<sub>4,97</sub> = 0.6850, p =0.6040<br>**P15 – P21 p < 0.0001**<br>**P15 – P25 p < 0.0001**<br>**P15 – P30 p < 0.0001**<br>**P18 – P21 p < 0.0001**<br>**P18 – P25 p < 0.0001**<br>**P18 – P30 p= 0.0004**<br>All other post-hocs p > 0.05. | P18 M: 11<br>P18 F: 8<br>P21 M: 12<br>P21 F: 13<br>P25 M: 12<br>P25 F: 10<br>P30 M: 11<br>P30 F: 11 |
| Fig. S3C (Spike number) | Paired t-test | **Light ON vs. light OFF: t<sub>5</sub> = 8.032, p = 0.0005** | 6 cells, 2 mice |
| Fig. S3D (RMP) | Paired t-test | **Light ON vs. light OFF: t<sub>5</sub> = 3.728, p = 0.0136** | 6 cells, 2 mice |
| Fig. S4B (Fear Training) | Two-way RM ANOVA | **Tone: F<sub>6,102</sub> = 36.45, p < 0.0001**<br>Sex: F<sub>1,17</sub> = 0.2456, p = 0.6265<br>Tone x sex: F<sub>6,102</sub> = 0.5279, p = 0.7859 | Male: 10<br>Female: 9 |
| Fig. S4C (Fear Training avg. tone) | Unpaired t-test | Male vs female: t<sub>17</sub> = 0.4741, p = 0.6415 | Male: 10<br>Female: 9 |
| Fig. S4D (Ext.) | Two-way RM ANOVA | **Tone: F<sub>1,17</sub> = 73.24, p < 0.0001**<br>Sex: F<sub>1,17</sub> = 0.3683, p = 0.5519<br>Tone x sex: F<sub>1,17</sub> = 0.1261, p = 0.7269 | Male: 10<br>Female: 9 |
| Fig. S4E (Ext. first 2 tones) | Unpaired t-test | Male vs female: t<sub>17</sub> = 0.5617, p = 0.5816 | Male: 10<br>Female: 9 |
| Fig. S4F (Ext. last 2 tones) | Unpaired t-test | Male vs female: t<sub>17</sub> = 0.1822, p = 0.8576 | Male: 10<br>Female: 9 |
| Fig. S4H (Fear Training) | Two-way RM ANOVA<br>Sidak’s multiple comparisons (collapsed sexes) | **Tone: F<sub>6,156</sub> = 56.18, p < 0.0001,**<br>**Age: F<sub>1,26</sub> = 31.24, p < 0.0001,**<br>Sex: F<sub>1,26</sub> = 1.573, p = 0.2209,<br>**Tone x age: F<sub>6,156</sub> = 5.777, p < 0.0001,**<br>Tone x sex: F<sub>6,156</sub> = 0.9235, p = 0.4798<br>Age x sex: F<sub>1,26</sub> = 2.411, p = 0.1326<br>Tone x age x sex: F<sub>6,156</sub> = 0.7137, p = 0.6391<br>Baseline: P21 – P30 p = 0.3982<br>Tone 1: P21 – P30 p = 0.2226<br>**Tone 2: P21 – P30 p < 0.0001**<br>**Tone 3: P21 – P30 p < 0.0001**<br>**Tone 4: P21 – P30 p = 0.0021**<br>Tone 5: P21 – P30 p = 0.2924<br>Tone 6: P21 – P30 p = 0.9996 | P21 M: 8<br>P21 F: 8<br>P30 M: 7<br>P30 F: 7 |
| Fig. S4I (Fear Training avg. tone) | Two-way ANOVA | **Age: F<sub>1,26</sub> = 33.86, p < 0.0001**<br>Sex: F<sub>1,26</sub> = 0.8486, p = 0.3654<br>Age x sex: F<sub>1,26</sub> = 0.8380, p = 0.3684 | P21 M: 8<br>P21 F: 8<br>P30 M: 7<br>P30 F: 7 |
| Fig. S4J (Ext.) | Three-way RM ANOVA | **Tone: F<sub>12,312</sub> = 9.078, p < 0.0001**<br>**Age F<sub>1,26</sub> = 10.18, p = 0.0037**<br>Sex: F<sub>1,26</sub> = 1.083, p = 0.3075<br>Tone x age: F<sub>12,312</sub> = 1.504, p = 0.1211<br>Tone x sex: F<sub>12,312</sub> = 1.749, p = 0.0561<br>Age x sex: F<sub>1,26</sub> = 1.464, p = 0.2371<br>Tone x age x sex: F<sub>12,312</sub> = 0.8478, p = 0.6009 | P21 M: 8<br>P21 F: 8<br>P30 M: 7<br>P30 F: 7 |
| Fig. S4K (Ext. first 2 tones) | Two-way ANOVA | **Age: F<sub>1,26</sub> = 17.18, p = 0.0003**<br>Sex: F<sub>1,26</sub> = 0.1744, p = 0.6796<br>Age x sex: F<sub>1,26</sub> = 1.220, p = 0.2795 | P21 M: 8<br>P21 F: 8<br>P30 M: 7<br>P30 F: 7 |
| Fig. S4L (Ext. last 2 tones) | Two-way ANOVA<br>Sidak’s multiple comparison’s test | Sex: F<sub>1,26</sub> = 0.0708, p = 0.7923<br>Age: F<sub>1,26</sub> = 1.142, p = 0.2951<br>**Age x sex: F<sub>1,26</sub> = 5.368, p = 0.0286**<br>**P21 – P30 males: p = 0.0478**<br>P21 – P30 females: p = 0.6223 | P21 M: 8<br>P21 F: 8<br>P30 M: 7<br>P30 F: 7 |
| Fig. S7B (Fear Training) | Two-way RM ANOVA | **Tone: F<sub>3.216, 35.37</sub> = 8.262, p = 0.0002**<br>Age: F<sub>1,11</sub> = 0.00316, p = 0.9561<br>Tone x age: F<sub>6,66</sub> = 0.4629, p = 0.8333 | P18 F: 7<br>P21 F: 6 |
| Fig. S7C (Fear Training avg tone) | Unpaired t-test | P18 vs P21: t<sub>11</sub> = 0.1393, p = 0.8917 | P18 F: 7<br>P21 F: 6 |
| Fig. S7D (Ext.) | Two-way RM ANOVA | **Age: F<sub>1,11</sub> = 6.179, p = 0.0303**<br>**Tone: F<sub>5.667, 62.34</sub> = 18.74, p < 0.0001**<br>Interaction: F<sub>12,132</sub> = 1.439, p = 0.1561 | P18 F: 7<br>P21 F: 6 |
| Fig. S7E (Ext. first 2 tones) | Unpaired t-test | P18 vs P21: t<sub>11</sub> = 1.489, p = 0.1646 | P18 F: 7<br>P21 F: 6 |
| Fig. S7F (Ext. last 2 tones) | Unpaired t-test | P18 vs P21: t<sub>11</sub> = 0.5274, p = 0.6084 | P18 F: 7<br>P21 F: 6 |
| Fig. S9C (AMPA: NMDA) | Unpaired t-test | **Control vs LTP: t<sub>7</sub>=3.132, p= 0.016** | 9 cells, 5 mice |

## Notes

### Competing Interest Statement

The authors have declared no competing interest.

